# Virulence adaptation of *Pseudomonas aeruginosa* phospholipase mutant with altered membrane phospholipid composition

**DOI:** 10.1101/2022.11.25.517918

**Authors:** Muttalip Caliskan, Gereon Poschmann, Mirja Gudzuhn, Daniel Waldera-Lupa, Wolfgang R. Streit, Karl-Erich Jaeger, Kai Stühler, Filip Kovacic

## Abstract

Membrane protein and phospholipid (PL) composition changes in response to environmental cues and during infections. Covalent modification and remodelling of the acyl chain length of PLs is an important bacterial adaptation mechanism. However, little is known about which bacterial pathways are regulated in response to altered PL composition. Here, we showed that *P. aeruginosa* phospholipase A, PlaF, which modulates membrane PL composition, is important for biofilm biogenesis, and we performed whole-cell quantitative proteomics of *P. aeruginosa* wild-type and Δ*plaF* biofilms to identify pathways regulated by PlaF. The results revealed profound alterations in the abundance of several two-component systems (TCSs), including accumulation of PprAB, which controls the transition to biofilm. Furthermore, a unique phosphorylation pattern of transcriptional regulators, transporters and metabolic enzymes, as well as differential production of seven proteases, in Δ*plaF*, indicate that PlaF-mediated virulence adaptation involves complex transcriptional and posttranscriptional regulation. Moreover, proteomics revealed the depletion of pyoverdine-mediated iron uptake pathway proteins in Δ*plaF*, which agrees with the decreased concentrations of extracellular pyoverdine and intracellular iron and is likely responsible for its prolonged lag growth phase, presumably due to reduced iron uptake. Conversely, the accumulation of proteins from alternative iron-uptake systems in Δ*plaF* suggests that PlaF may function as a switch between different iron-acquisition pathways. The observation that Δ*plaF* accumulates PL-acyl chain modifying and PL synthesis enzymes reveals novel insights into the role of PlaF for membrane PL homeostasis. Although the precise mechanism by which PlaF simultaneously affects multiple pathways remains to be elucidated, we suggest that PlaF-catalyses the degradation of PLs which then serve as a signal that is amplified by proteins of two-component, phosphorylation and proteolytic degradation systems to elicit the global adaptive response in *P. aeruginosa*.

## Introduction

The full biological activity of membrane proteins is ensured through their interactions with lipids and proteins. Protein-lipid interactions regulate protein function and are responsible for the temporal and spatial organization of membrane complexes and functional membrane microdomains [1–5]. An increasing amount of structural and biochemical data has revealed functional modulation of integral membrane proteins through interactions with specific phospholipids (PLs), for example, the *E. coli* magnesium transporter MgtA [6], the SecYEG protein secretion translocon [7] and the anaerobic respiratory complex NarGHI of *E. coli* [8]. Interactions between some integral membrane proteins and specific PLs assist protein folding, resulting in protein activation, as described for lactose permease LacY of *E. coli* [9], or activity switching, as described for the fatty acid desaturase DesK of *Bacillus subtilis* [10]. The binding of soluble proteins to the membrane and their allosteric activation, known as interfacial activation, was described to be lipid dependent for human phospholipase A2 [11] and 5-lipooxygenase [12] and bacterial ATPase SecA [7, 13]. In addition, phospholipids have been demonstrated to facilitate homodimerization of SecYEG [14, 15] and succinate dehydrogenase [16], hetero-oligomerization of the cytochrome *bc*_1_ complex (complex III) [17] and the barrel assembly machine (Bam) complex [18, 19], and even the assembly of multisubunit complexes in respiratory supercomplex III_2_IV [20] and photosystem II-light-harvesting complexes II supercomplex [21].

As a consequence of the regulation of membrane proteins by lipids, bacteria tailor membrane PL compositions to endow cellular membranes with defined properties. This is apparent from recent lipidomic analyses that reveal the adaptation of the chemical composition of bacterial membrane PLs to numerous environmental and developmental conditions, including planktonic growth [22], biofilm [22], hyperosmotic stress [23], stationary growth phase [24], iron limitation [25], increased temperature [26], pH stress [27], and antibiotic stress [28, 29]. Although these conditions exert changes on the bacterial proteome [30–36], it remains elusive which pathways are lipid-dependent and how bacteria simultaneously modulate their lipidome and proteome under dynamic conditions.

Recently, we discovered two intracellular PLAs, PlaF and PlaB, in *P. aeruginosa* with the ability to hydrolyse endogenous membrane PLs [37, 38]. *In vivo* PlaF degradation of six PLs differing in head groups and acyl chains was suggested to be responsible for alterations of their membrane PL composition [38] through a pathway analogous to the eukaryotic Lands cycle PL remodelling pathway [39]. PL remodelling mechanism suggested for PlaF, is similar to the recently discovered *P. aeruginosa* pathway involving a phospholipase C and glycosyltransferase-catalysed replacement of glycerophospholipids of outer membrane with glycolipids in response to phosphate limitation [40]. The function of PlaF in *in vivo* PL modulation was strengthened by findings that PlaF shows *in vitro* catalytic activity towards physiological substrates and that PlaF activity is regulated by a negative feedback loop based on an association with a reaction product, fatty acids (FAs) [38].

A crystal structure and biochemical and computational results provided a mechanism of how dimer to monomer transition of PlaF and subsequent rotation of PlaF at the membrane surface leads to recognition of PL substrate, its extraction from the membrane through the tunnel to the active site where it is hydrolysed and the products are likely released to the membrane [38, 41, 42]. We suggest that PlaF-dependent modulation of the membrane PL composition is responsible for the strongly attenuated virulence of *P. aeruginosa ΔplaF* in a *Galleria mellonella* virulence assay and macrophage cytotoxicity assay.

This is in agreement with results showing that alteration of the PL composition affects the virulence properties of *P. aeruginosa* [22, 28, 43] and other human pathogens [25, 44].

Here, we used *P. aeruginosa ΔplaF* to identify pathways depending on PlaF-mediated remodelling of the PL composition. Whole-cell quantitative proteomics revealed an unprecedented molecular view of the relationship between *P. aeruginosa* membrane PL composition and sensing and signal transduction, protein phosphorylation and degradation and PL homeostasis pathways. Phenotypic characterization confirmed that the PlaF-dependent alteration of multiple pathways in *P. aeruginosa ΔplaF* regulates biofilm biogenesis, cell motility and iron acquisition.

## Results

### Dysfunctional biofilm formation in P. aeruginosa ΔplaF is related to global proteomic changes

*P. aeruginosa* alters its lipidome upon transition to biofilm life style and attachment to surfaces [22]; therefore, we analysed whether *P. aeruginosa* Δ*plaF* (in following text designated as Δ*plaF*) with an altered lipidome would affect biofilm. To do so, we performed comprehensive biofilm studies of *P. aeruginosa* WT and Δ*plaF* grown in a microtiter plate (MTP) under static conditions and in a flow chamber constantly supplemented with nutrients. We observed that *P. aeruginosa ΔplaF* grown under static conditions showed a significant (*n* = 40, *p* < 0.05) reduction (15–72%) in the amount of biofilm produced compared with the wild-type strain after 8, 24, and 72 h of growth (Fig. 1A). No significant difference was observed after 6 and 9 days of growth in MTPs (Fig. 1A).

**Figure 1:**
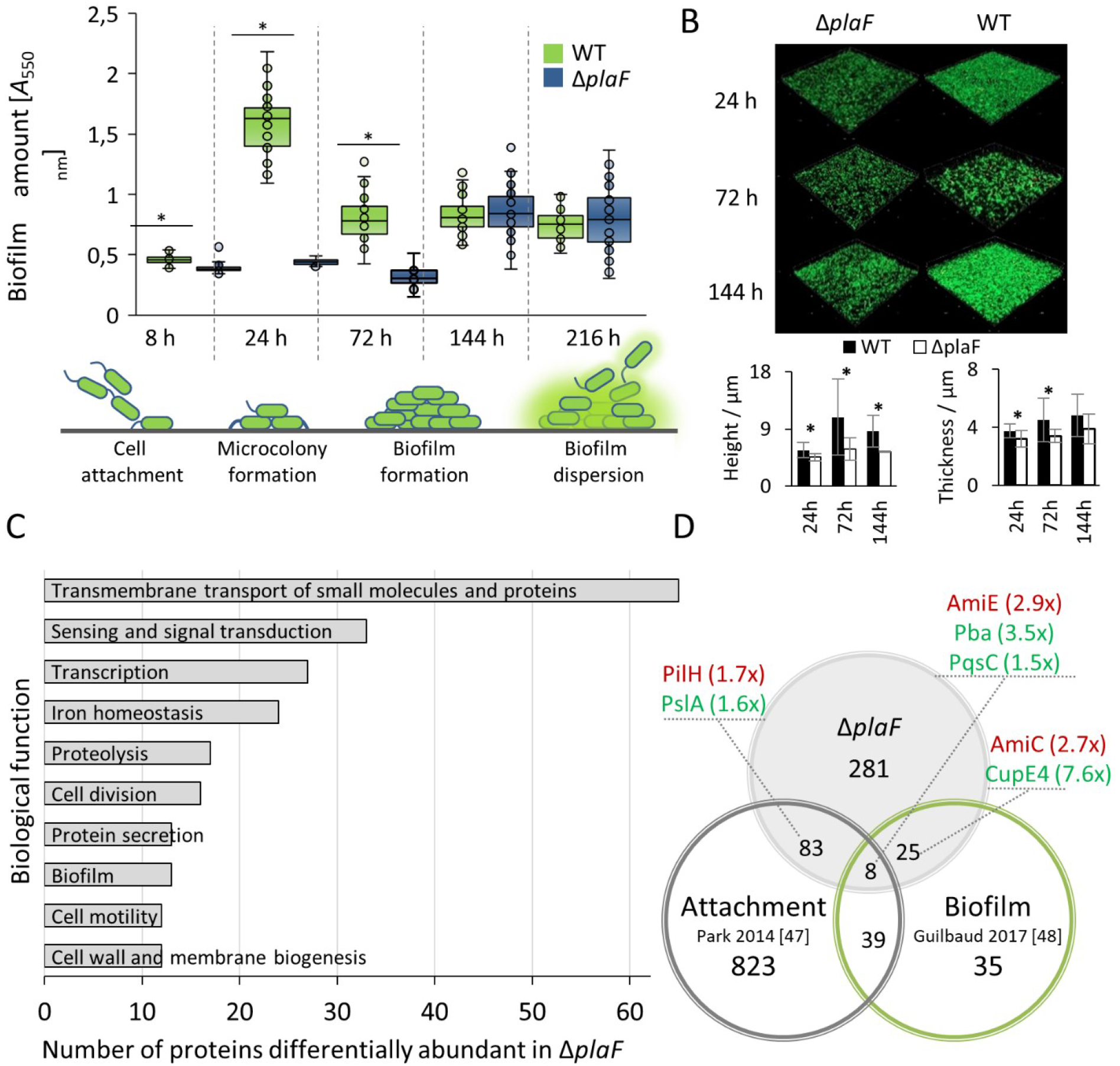
PlaF is a global regulator of *P. aeruginosa* biofilm. **A)** Amount of biofilm produced by the *P. aeruginosa ΔplaF* and WT strains cultivated in 96-well MTPs (LB medium, 37 °C, without aeration) was quantified by staining the cells attached to the MTP with crystal violet. The results are the mean ± standard deviation of five biological replicates, each measured eight times. Statistical analysis was performed using Student’s *t* tests, * p < 0.05. **B)** Biofilm architecture was analysed by CLSM after 24, 72, and 144 h of growth at 37 °C in a flow cell with a continuous supply of LB medium. Representative figures of two biological replicates analysed by imaging 100 × 100 μm sections of the cover glass. More CLSM images are shown in Fig. S1. Biofilms were quantified by analysing 8 pictures per condition using BiofilmQ software; * p < 0.05. **C)** Functional classification of proteins differentially abundant in WT and Δ*plaF*. Functional assignment according to biological processes was performed with COG [46]. The GO identifiers of each functional group are listed in Table S4. **D)** Overlap of proteins identified as differentially abundant in our study and proteins identified as biofilm- [47] and attachment-related [48] in proteomic studies by other researchers. Red and green indicate proteins less and more abundant in Δ*plaF*, and numbers in brackets show a fold change.

The CLSM results of biofilms grown for 24, 72, and 144 h on glass cover slides revealed considerable differences in the architecture between *P. aeruginosa ΔplaF* and WT (Figs. 1B and S1). The WT biofilm nearly homogeneously and densely covered the glass surface after 24 h, in contrast to that for Δ*plaF*, which showed a less dense and inhomogeneous structure. After 72 h, the biofilm of the WT strain consisted of large cell aggregates, while small aggregates were observed for the Δ*plaF* biofilm. After six days, the number of cell aggregates in the Δ*plaF* biofilm increased nearly to the level of the WT biofilm. Quantification of biofilms revealed significant (*p* < 0.05) different height between the WT and Δ*plaF* during 144 h while mean thickness was significantly different during 72 h.

These results show that the early phase of attachment to plastic and glass surfaces and biofilm maturation are negatively affected by PlaF. To understand the underlying molecular mechanisms involved in PlaF-mediated regulation of biofilms and their previously observed effects on virulence [38], we performed a whole-cell comparative-proteomic study of Δ*plaF* and WT grown under static biofilm conditions. Using electrospray-ionization mass spectrometry, we identified 2952 proteins, of which 2218 were accurately quantified in the two strains (Table S1, Figs. S2 and S3). The statistical analysis of five biological replicates using stringent criteria (number of unique peptides ≥ 2, *p* ≤ 0.05) revealed 397 proteins with an abundance ≥ 1.5-fold (120 proteins showed ≥ 2.0-fold change). Among them, 211 proteins were more abundant and 186 were less abundant in Δ*plaF* (Table S2). We observed profound alteration of the cell envelope, including the cytoplasmic membrane, which accommodates PlaF. This was indicated by one-dimensional annotation enrichment scores [45] for the inner membrane, periplasm and outer membrane of 0.30, −0.50 and −0.46 (Benjamini Hochberg corrected *p* values, 3.6 × 10^-5^ - 6.7 × 10^-10^) (Table S3). Interestingly, we also observed a significant (*p* = 0.001) alteration of the abundance of extracellular proteins in Δ*plaF*, which indicates an effect of PlaF on protein secretion and/or extracellular matrix remodelling.

The functional classification of differentially abundant proteins, based on the Cluster of Orthologous Groups (COG) database [46], revealed that the majority have membrane-associated functions, e.g., transmembrane transport of small molecules and proteins (65 proteins), sensing and signal transduction (33 proteins) and cell wall and membrane biogenesis (12 proteins) (Fig. 1C, Table S4). Furthermore, PlaF exerts an effect on various important cellular processes, including transcription (27 proteins), iron homeostasis (24 proteins), proteolysis (17 proteins) and cell division (16 proteins). Not surprisingly, many differentially abundant proteins were related to biofilm biogenesis (13 proteins) and other virulence traits, e.g., protein secretion (13 proteins) and cell motility (12 proteins) (Fig. 1C).

The comparison of proteins differentially abundant in Δ*plaF* with the proteomic datasets of experimentally identified proteins involved in attachment to glass and plastic surfaces [48] and biofilm formation [47] revealed 97 and 15 shared proteins, respectively (Fig. 1D, Table S5). Among them were several proteins whose roles in attachment and biofilm formation are well understood. For example, PilH which regulates the function of type IV pili important for sensing solid surfaces [49], and the CupE4 chaperone essential for assembling cell surface fimbriae structures and promoting biofilm formation [50]. Furthermore, we observed a much higher abundance (+3.5-fold) of an outer membrane protein (Pba) in Δ*plaF* (Fig. 1D). This protein was recently demonstrated to function in biofilm organization by binding to the peptidoglycan layer [51]. Quantitative proteomic studies of two other research groups showed that Pba is a biofilm-specific protein [47] and that *P. aeruginosa* produces less Pba upon adherence to the surface [48]. This result agrees with our results, which reveal impaired adherence to the plastic surfaces and overproduction of Pba by Δ*plaF* (Fig. 1B).

Through the synthesis of *Pseudomonas* quinolone signal (PQS), PqsC regulates quorum-sensing signalling cascades involved in attachment and biofilm formation [52, 53]. Our results showed that PqsC was slightly more abundant in Δ*plaF* (+1.5-fold), so we analysed the relationship between PlaF and the QS network. We did not observe the overproduction of other enzymes from the PQS synthesis pathway, as PqsD, PqsH, and PqsH did not significantly accumulate in Δ*plaF* and PqsB and PqsA showed only +1.4-fold and +2.3-fold higher abundance in Δ*plaF*. Furthermore, among the 397 PlaF-affected proteins presented here, only 22 were identified as QS-regulfseeated [54], and only two of them, BphP phytochrome and putative hybrid two-component protein PA0178, showed > 2-fold changed protein abundance in Δ*plaF* (Fig. S4). This result, together with the finding that PlaF showed only negligible activation (1.4-fold) in a comprehensive transcriptomics analysis of a QS regulon by Asfahl *et al*. [54], suggests that PlaF affects the biofilm formation and virulence of *P. aeruginosa* in a PQS-independent manner.

The second most differentially abundant protein was the two-component sensor PprA (+18.7-fold in Δ*plaF*), which is a key regulator of the transition between planktonic and biofilm lifestyles [55]. Another putative two-component sensor, PA0178, was also strongly (+4.6-fold) affected in Δ*plaF*. Furthermore, our results revealed the most substantial negative effect of PlaF on the abundance of iron homeostasis proteins from the pyoverdine pathway, as PvdT, PvdN, PvdM, PtaA and PvdR showed between 6.1 and 3.1 less abundance in Δ*plaF* than in WT. The interdependence between two-component system-mediated sensing and transduction, biofilm formation and iron acquisition is well described in *P. aeruginosa* [56, 57]; therefore, we studied the role of PlaF in these processes in more details.

### PlaF alters TCS-mediated virulence adaptation, including the PprAB-controlled switch from a planktonic to a biofilm lifestyle

Of approximately 65 TCSs coordinating virulence adaptation and biofilm formation in *P. aeruginosa* PA01 [57], 14 showed significantly different protein abundance between Δ*plaF* and WT. For six of them, only one component, sensor kinase (SK) or response regulator (RR), could be identified (Fig. 2A). It is notable that the large majority of TCS proteins accumulated in Δ*plaF*, including four SK, seven RR, and two hybrid TCS (Fig. 2A). For most of them, the signal triggering the response is unknown, while for several of them, it was reported that they sense extracellular phosphate (PhoR) [58], iron (PmrAB) [59], potassium (KdpD) and tricarboxylic acids (TctD) [60]. Not surprisingly, many affected TCS, including RocA1, PhoR, FleR, PA4396, TctD, PprA and PprB, are involved in the regulation of biofilm formation and adhesion [57, 61]. This massive imbalance of TCSs in Δ*plaF* may affect the transcriptional regulation of biofilm biogenesis genes, thereby contributing to the reduced biofilm formation of this strain.

**Figure 2:**
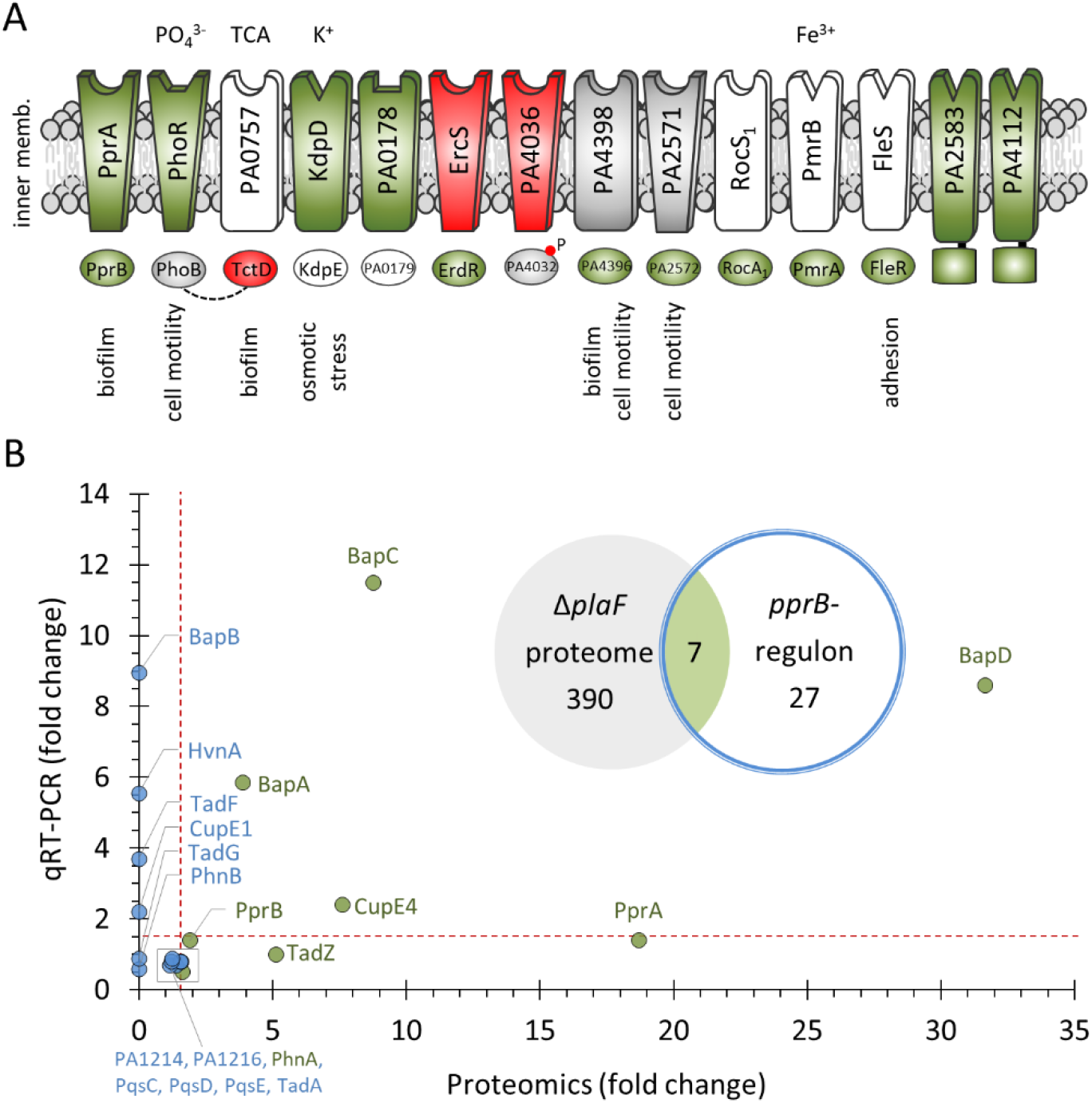
Two-component system sensing and signal transduction are affected in Δ*plaF*. **A)** TCS proteins depleted, accumulated, unaffected and not identified in Δ*plaF* are indicated as red, green, grey and white, respectively. The signalling molecules and regulated cellular processes are indicated above and below the respective TCS. The dashed line indicates the functional interaction of RRs and the red circle (P) denotes phosphorylation. **B)** The correlation between protein and mRNA (determined with qRT–PCR) abundance for biofilm-related proteins/genes in Δ*plaF*. Red dashed lines indicate the threshold of 1.5-fold change. Inset: overlap of differentially abundant proteins identified in the Δ*plaF* proteome (our study) with the differentially transcribed genes in *P. aeruginosa ΔpprB* [55].

The most severely affected TCS was PprAB, in which SK was +18.7-fold and RR was +1.9-fold more abundant in Δ*plaF*. This result was expected, knowing that PprAB TCS regulates the transition from planktonic to biofilm lifestyles, although the signal sensed by PprA is not yet known [50, 62]. However, less biofilm production by Δ*plaF* contrasts with the finding that overexpression of the PprAB TCS enhances biofilm formation [55]. To better understand this phenomenon, we compared our proteomics results with the results of a transcriptomics study that identified 34 genes of *P. aeruginosa* that were transcriptionally regulated by *pprB* [62]. The proteomics results agreed with the transcriptomics results, which revealed that overexpression of the *pprB* gene (32.2-fold) upregulates transcription of the *pprA* gene (11.5-fold); however, at the protein level, the effect was much stronger, as 1.9-fold overproduction of PprB resulted in nearly 19-fold overproduction of PprA. Such a different PprA/PprB ratio in transcriptomics (0.3) and proteomics (10.0) studies and the fact that other biofilm-related TCSs are affected in Δ*plaF* may be an explanation for the unexpected biofilm phenotype observed in Δ*plaF*.

Proteomics analysis identified 12 proteins encoded by genes that are transcriptionally regulated by the PprAB TCS. All of them accumulated in Δ*plaF*, yet only seven showed significant differences (Fig. 2B, inset), which was expected, as PprB was more abundant in Δ*plaF*. The biofilm-associated protein BapD belonging to the *pprAB* regulon was, in our proteomic analysis, the most highly abundant (+31.7-fold in Δ*plaF*) among the entire set of PlaF-affected proteins. BapD is an adhesin secreted by an ABC efflux pump encoded by BapA, BapB and BapC; all four are organized in an operon [63]. Approximately a 9- and 4-fold higher abundancies of the ABC transporter proteins BapC and BapA were observed in Δ*plaF*, respectively, while the outer membrane protein BapB could not be identified in our proteomics study, probably because of its hydrophobicity. Furthermore, two other PprAB-regulated proteins, CupE4 (+7.6-fold) and TadZ (+5.1-fold), strongly accumulated in Δ*plaF*. The chaperone/usher pathway protein CupE4 and the tight adherence protein TadZ, together with other Cup and Tad subunits, form fimbrial structures on the cell surface, which contribute to the biofilm formation of *P. aeruginosa* [55]. For unknown reasons, our proteomic study did not detect other fimbrial proteins of the Tad and Cup complexes.

To better understand PprAB-mediated gene regulation in Δ*plaF*, we investigated this regulatory network at the transcriptional level using a quantitative real-time-PCR (qRT-PCR) approach. The results of qRT-PCR revealed upregulation of *bapC, bapD*, and *cupE4*, which were also more abundant in Δ*plaF* at the protein level, and *bapB, cupE1, hvnA*, and *tadF*, whose products were not identified by proteomics. TadA, PqsC, PqsD, PqsE, PA1214, and PA1216, which did not show differential protein abundance in Δ*plaF*, and TadG and PhnB, which were not identified by proteomics, were also not upregulated in Δ*plaF*, as determined by qRT–PCR (Fig. 2B). These results show that PprAB regulated gene transcription is upregulated in Δ*plaF*, although the amount of PprB is likely insufficient for the activation of all genes required for proper biofilm formation, which may be one of the reasons for the lower biofilm formation observed in Δ*plaF*.

### *PlaF affects phosphorylation and proteolytic degradation pathways of* P. aeruginosa

TCS-regulated gene expression relies on the phosphorylation of RR by respective SK; therefore, we next analysed proteomics results to identify phosphorylated proteins in the Δ*plaF* and WT strains. In an approach that did not involve the enrichment of phosphopeptides, phosphorylated S, T or Y residues of nine proteins were identified, and phosphopeptides were quantified to obtain their ratio in Δ*plaF* compared to WT normalized to the total protein amount (Fig. 3A and Table S6). To our knowledge, only anti anti-sigma factor HsbA and isocitrate dehydrogenase were previously reported to be phosphorylated in *P. aeruginosa* [64–66].

**Figure 3:**
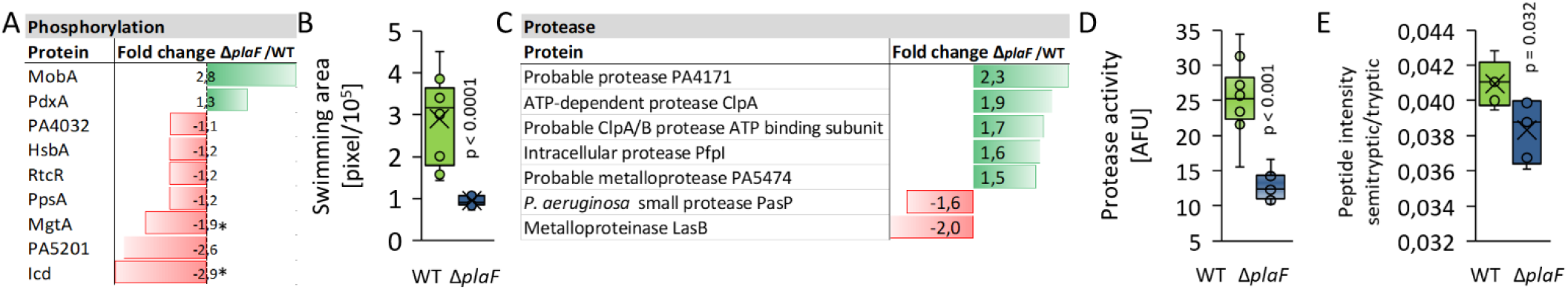
Phosphorylation, swimming and protein turnover systems are imbalanced in Δ*plaF*. **A)** Normalized abundance of phosphorylated peptides identified by MS analysis in at least 4 out of 5 biological replicates of WT and Δ*plaF*. Green and red bars indicate protein specific phosphopeptides accumulated and depleted in Δ*plaF*. Detailed sequences of phosphopeptides and data from quantification are shown in Table S6. The proteins indicated by * showed significant (*p* < 0.05) changes in phosphopeptide abundance. **B)** Swimming motility of Δ*plaF* is impaired in comparison to the WT. Growth after 24 h at 37 °C on the LB agar swimming plate was quantified using ImageJ and expressed as the swimming area. The assay was performed with 9 biological replicates. **C)** Proteases that are significantly (*p* < 0.05) differentially abundant in WT and Δ*plaF* as identified by MS analysis. Green and red bars indicate proteins accumulated and depleted in Δ*plaF*. **D)** Intracellular protease activity measured with the FTC-casein assay. **E)** Number of valid intensity values for semitryptic/tryptic peptides.

Interestingly, among the identified phosphoproteins, the phosphorylated peptide of only one putative TCS RR, PA4032, did not show significant differences in abundance between WT and Δ*plaF* (*p* = 0.91). The respective putative SK PA4036 was 1.5-fold less abundant in Δ*plaF*. Notably, the phosphorylation of the remaining nine RRs identified in the proteome could not be detected despite the accumulation of several SKs in Δ*plaF*. This result indicates that SKs accumulated in Δ*plaF* might have impaired kinase function or enhanced phosphatase function. It remains to be answered whether the function of the SKs identified here could be regulated by PlaF-mediated alteration of PLs, as recently described for lipid kinases [67].

The normalized abundance of phosphopeptides of P-type ATPase MgtA (−1.9-fold) involved in citrate-mediated iron uptake was significantly (*p* < 0.02) lower in Δ*plaF*. During each cycle of metal ion transport, MgtA is activated by autophosphorylation followed by dephosphorylation to regenerate the transporter for a new cycle [68]. In *E. coli*, MgtA phosphorylation of residue D373 in the conserved DKTGT pentapeptide leads to activation of Mg^2+^ transport [6]. Although the DKTGT-pentapeptide is conserved in *P. aeruginosa* MgtA, we did not find phosphorylation within this motif. Our MS data revealed that phosphorylation of *P. aeruginosa* MgtA takes place on Y215 or T217, which are located in a cytoplasmic actuator domain. Hence, Y215 is substituted by Ala in *E. coli* MgtA, while T217 is conserved (Fig. S5). The actuator domain has an important dual function as it undergoes conformational changes upon D373 phosphorylation to activate *E. coli* MgtA, and it dephosphorylates D373 to inactivate the transporter [6]. Although the precise phosphorylation site of the actuator domain of *P. aeruginosa* MgtA and the functional consequences of phosphorylation remains to be elucidated, our results suggest differences between *E. coli* and *P. aeruginosa* MgtA, which may indicate their different functions in the transport of Mg^2+^ and the Fe^2+^:citrate complex into the cytoplasm [6, 69].

The normalized abundance of phosphopeptides of isocitrate dehydrogenase Icd from the tricarboxylic acid (TCA) cycle was significantly lower (−2.9-fold, *p* < 0.02) in Δ*plaF*. Phosphorylation of the serine residue at position 115, also detected in our study, was reported as a mechanism of Icd inactivation [64]; therefore, the accumulation of dephosphorylated Icd in Δ*plaF* indicates upregulation of Icd-catalysed oxidative decarboxylation of isocitrate in the TCA cycle [64]. The abundance of phosphatase AceK, responsible for Icd phosphorylation, was not significantly (*p* = 0.78) different in Δ*plaF*; therefore, it is likely that allosteric regulation of AceK by metabolites (acetyl-CoA and oxaloacetate) [64] may be responsible for the differential phosphorylation of Icd. In *P. aeruginosa*, Icd competes for isocitrate with isocitrate lyase (Icl), a key enzyme within the glyoxylate shunt (GS) [64]. Proteomics results revealed that Icl was slightly (−1.2-fold) yet significantly (*p* = 0.03) less abundant in Δ*plaF*. The increased amount of dephosphorylated Icd and decreased amount of Icl in Δ*plaF* indicates a downregulation of GS. GS is upregulated upon catabolism of FAs to acetyl-CoA [70], a building block of FAs and allosteric regulator of TCA cycle, therefore, the suggested downregulation of GS in Δ*plaF* agrees with the expected reduction in release of FAs from endogenous PLs in Δ*plaF* [38].

We have shown that the anti-anti-sigma factor HsbA, which is significantly accumulated (+1.5-fold, *p* < 0.01) in Δ*plaF*, is phosphorylated at S56. Note that the normalized abundance of HsbA-phosphopeptide in Δ*plaF* is not significantly different from that in the WT. This is in agreement with the observation that the kinase HsbR, an enzyme that catalyses the phosphorylation of HsbA [71], is similarly abundant in Δ*plaF* and WT. The phosphorylation of HsbA is crucial for its pivotal role in the regulation of a complex signalling network that controls cell motility and biofilm biogenesis [66, 71, 72]. Therefore, we tested motility and observed that swimming was significantly impaired in Δ*plaF* in contrast to swarming and twitching, which were not affected (Figs. 3B and S6). Furthermore, it was demonstrated that the HsbR-HsbA system [71] posttranslationally regulates the alternative sigma factor RpoS, which was +2.8-fold (*p* = 0.001) more abundant in Δ*plaF*. This may partially explain the reduced biofilm formation of Δ*plaF* because *P. aeruginosa* Δ*rpoS* was shown to overproduce biofilm compared to the wild-type strain [73].

Transcriptional regulation is counterbalanced by the action of proteases that regulate various pathways, including those involved in signalling and biofilm formation. Quantitative proteomic results revealed that several proteases (ClpA, PasP, PA4171, and PA5474) were significantly accumulated (to +2.3-fold) and several other proteases (LasB and PasP) were significantly depleted (to −2.0-fold) in Δ*plaF* (Fig. 3C). Therefore, we assayed the intracellular protease activity of Δ*plaF* and WT using fluorescently labelled casein, a general substrate used for assaying a broad range of proteases. The results revealed 50% (*n* = 10, *p* < 0.001) lower proteolytic activity in Δ*plaF* than in the WT (Fig. 3D). This result was tested by proteomic analysis of peptides generated from proteins partially degraded by host proteases, as these semitryptic peptides contain only one trypsin restriction site. Indeed, proteomics results confirmed significantly (*p* = 0.032) higher abundance of semitryptic peptides in the WT than in Δ*plaF* (Fig. 3E). Among 27 semitryptic peptides that showed significantly different abundance in Δ*plaF* were peptides from proteins that belong to PlaF-regulated pathways, iron homeostasis (MgtA, FpvA), biofilm formation (biofilm dispersion protein BdlA) and gene transcription (RpoS) (Table S7). Our previous results of tryptic peptide MS analysis showed that the abundance of MgtA, FpvA and RpoS differed in the WT and Δ*plaF*, while BdlA was not detected. This result indicates that the altered protein turnover system in Δ*plaF* accounts for partial and complete degradation of protease substrates, which may, in part, contribute to observed differences in the abundance of proteins in Δ*plaF*.

In conclusion, proteomics analysis of phosphorylated and semitryptic peptides in Δ*plaF* revealed unique phosphorylation and protein degradation patterns, indicating that the PlaF-mediated virulence adaptation in *P. aeruginosa* involves transcriptional and posttranscriptional regulation, which rely on phosphorylation and proteolysis cascades.

### Pyoverdine-mediated iron acquisition is downregulated and alternative iron uptake systems are upregulated in ΔplaF

For growth and host colonization, *P. aeruginosa* relies on iron, which is thus a reason for the development of multiple systems mediating iron transport into the cell [74]. By mapping our proteomics results onto a comprehensive *P. aeruginosa* iron-homeostasis (*Pa*FeHo) network derived from the literature (Table S8), we identified 26 significantly (*p* < 0.05) differentially abundant proteins in Δ*plaF* (Fig. 4A). Among the proteins depleted in Δ*plaF*, the strongest effect was observed for the proteins constituting the pathway facilitating iron uptake by pyoverdine, a major iron-binding chelator in *P. aeruginosa*, [75, 76] (Fig. 4B). Five cytoplasmic proteins responsible for the synthesis of ferribactin, a precursor of pyoverdine, as well as almost all proteins involved in the periplasmic maturation of ferribactin to pyoverdine, were less abundant in Δ*plaF*. Proteins mediating the transport of ferribactin (PvdE) and pyoverdine (PvdRT) across the cytoplasmic and outer membranes, respectively, also showed lower abundance. Furthermore, in Δ*plaF*, FpvA, an outer membrane receptor for the pyoverdine:Fe^3+^ (Pvd:Fe^3+^) complex; TonB1, which provides energy to FpvA for active transport of Pvd:Fe^3+^ across the outer membrane; and FpvF and FpvC, whose functions are related to periplasmic sensing and release of Fe^3+^ from Pvd:Fe^3+^, were depleted. Moreover, the cytoplasmic membrane Fe^3+^ transporter HitA [77] was depleted in Δ*plaF*, although its functions in pyoverdine-mediated iron acquisition are not entirely understood [69]. A significant decrease in extracellular pyoverdine by approximately 40% in Δ*plaF* biofilms compared to WT biofilms strongly suggests a role for PlaF in the regulation of the pyoverdine pathway for iron utilization (Fig. 4C). A similar effect was observed when Δ*plaF* was grown under nonadherent conditions (Fig. 4C).

**Figure 4:**
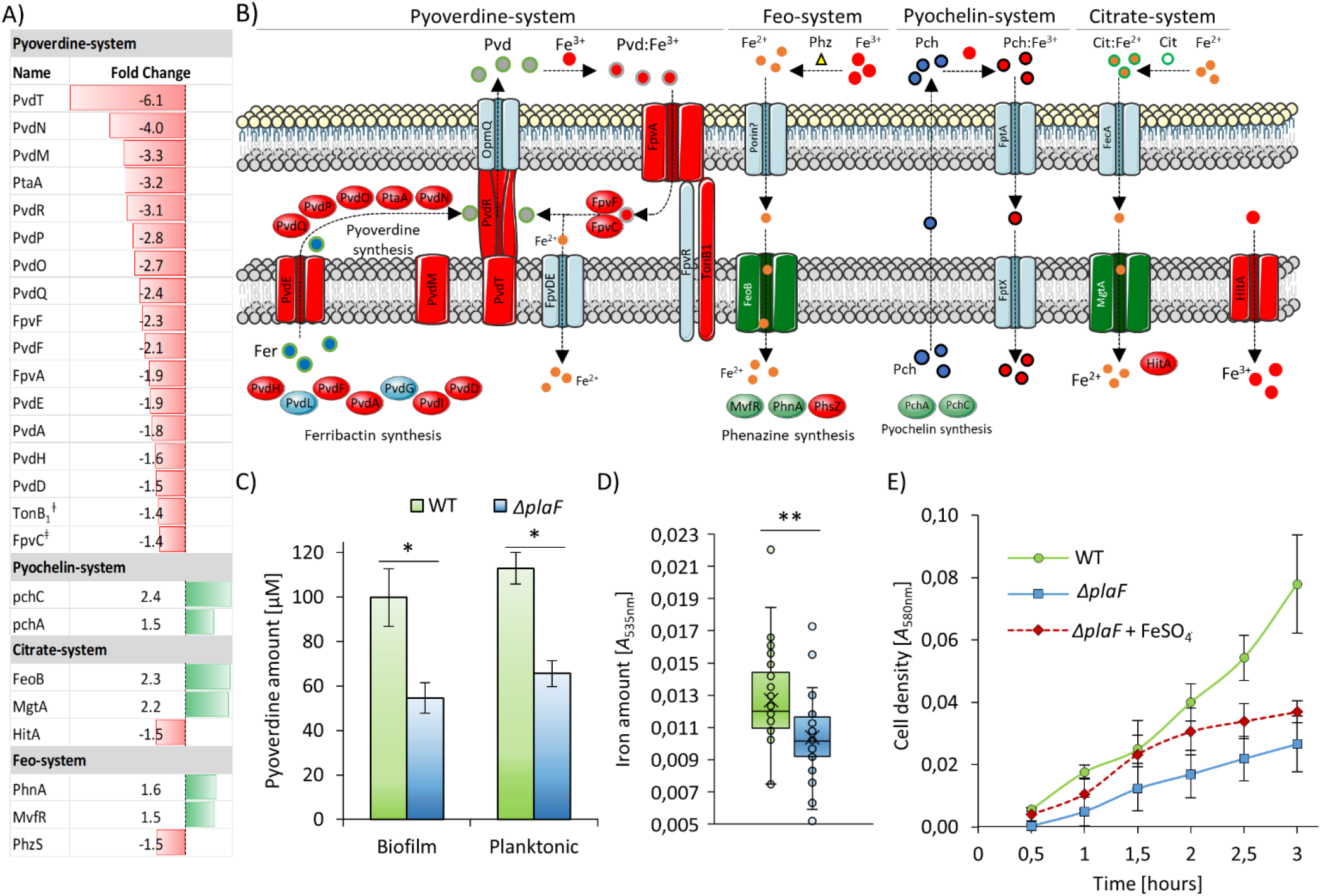
Iron acquisition is imbalanced in *P. aeruginosa ΔplaF*. **A)** Iron homeostasis proteins identified by proteomics analysis showing higher (green bars) and lower (red bars) abundance in Δ*plaF*. ÷FpvC and TonB1 abundance was slightly below the cut-off of 1.5-fold. **B)** The proteins for pyoverdine-, pyochelin- and citrate-mediated iron acquisition, as well as for Fe^3+^ and ferrous iron transport [69, 76] in *P. aeruginosa ΔplaF*. Proteins that were less and more abundant in *P. aeruginosa ΔplaF*are indicated in red and green, respectively. The blue-coloured proteins were not affected or not identified in our proteomics study. **C)** Pyoverdine concentration was quantified in the cell-free supernatant of WT and Δ*plaF* (5 biological replicates) grown in plastic MTPs (biofilm) or Erlenmeyer flasks (planktonic) at 37 °C for 24 h. The results are the mean ± SD. **D)** The intracellular iron concentration of 24 advanced biofilms was determined using 10 biological replicates. The results are the mean ± SD. **E)** The growth curves of WT and Δ*plaF* (3 biological replicates) in M9 minimal medium under iron-limiting conditions and in the presence of 100 μM FeSO_4_ were determined by measuring the optical density at 580 nm. The results are the mean ± SD. The growth of Δ*plaF* and WT, as well as Δ*plaF* and Δ*plaF* + FeSO_4_, after 2.5 h and 3 h differed significantly (*p* < 0.05).

We further analysed the abundance of the sigma factors PvdS and FpvI, which control the transcription of the Pvd and Fpv genes of the pyoverdine pathway, respectively. However, they were not found among the proteins quantified in our proteomics analysis, although the Fur protein, which regulates the expression of PvdS and FpvI, was identified as similarly abundant in both analysed strains [78]. Presently, the mechanism by which PlaF exerts an effect on the pyoverdine pathway remains unknown, and further studies are needed.

Conversely, many proteins involved in pyoverdine-independent iron uptake pathways accumulated in Δ*plaF* (Fig. 4B). Among them are PchA and PchC, which are involved in the synthesis of pyochelin, the second major iron-chelating siderophore; FeoB, PhnA and MvfR, which belong to the Feo-pathway and are responsible for the uptake of ferrous iron using phenazines [79, 80]; and the citrate:Fe^2+^ transporter MgtA [69]. Apparently, the deletion of the *plaF* gene in *P. aeruginosa* leads to a lower abundance of pyoverdine-mediated iron uptake proteins and cellular responses involving an increase in the abundance of proteins of alternative iron-acquisition systems.

Such feedback regulation may ensure to maintain a physiological level of intracellular iron. Iron is an essential metal ion that is accumulated by the bacteria in the lag phase to be used as a co-factor of primary metabolism enzyme during growth. [81, 82]. Therefore, we investigated the growth of the WT and Δ*plaF* in chemically defined iron-depleted medium. The results revealed significantly slower growth of Δ*plaF* during lag phase (0-3 h); which was not observed after 24 h (Figs. 4D and S7). The initial growth of Δ*plaF* was only slightly increased by supplementation with 100 μM FeSO_4_ in the medium (Fig. 4D). These results indicate that PlaF might play a role in acquisition of iron during the lag phase and the adaptation of *P. aeruginosa* to switch between iron uptake systems.

### *Phospholipid homeostasis is affected in* P. aeruginosa Δ*plaF*

*P. aeruginosa* biofilms show differences in the lipidome compared to planktonic cells [22, 38]. As PlaF affects biofilms and accounts for changes in the PL content of *P. aeruginosa* membranes [38], we analysed the proteomics results to understand the relationship between PlaF and PL homeostasis. For this purpose, Δ*plaF* proteomic results were mapped onto a comprehensive *P. aeruginosa* PL-homeostasis (*Pa*PLHo) network derived from the literature (Table S9 and Fig. S8). Among 38 proteins in the *Pa*PLHo network (Table S9), the cyclopropane fatty acid synthase Cfa (+1.8-fold), cis-trans isomerase Cti (+1.6-fold) and phosphotransacylase PlsX (+1.5-fold) were significantly accumulated in Δ*plaF* (Fig. 5A). The higher abundance of Cfa in Δ*plaF* is in agreement with the accumulation of its transcriptional activator RpoS [83].

**Figure 5:**
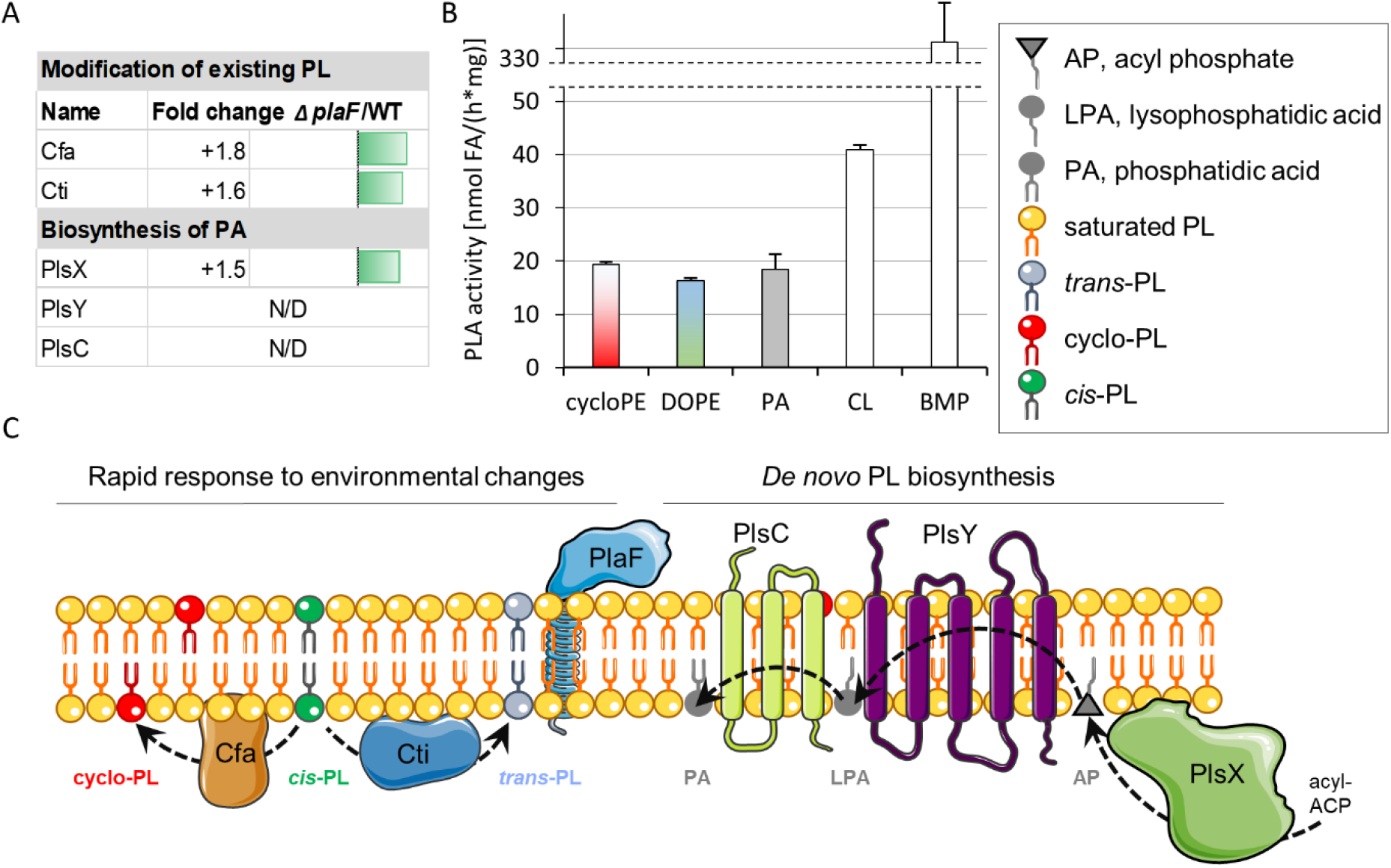
Phospholipid homeostasis is affected in *P. aeruginosa ΔplaF*. A) PL homeostasis proteins significantly (*p* < 0.05) more abundant in Δ*plaF* identified by proteomics analysis. Cti, *cis-trans* isomerase; Cfa, cyclopropane fatty acid synthase. The lysophosphatidic acid synthase PlsY and phosphatidic acid synthase PlsC were not detected in our proteomics study. B) *In vitro* PLA activity of PlaF with cyclopropyl phosphatidylethanolamine (cyclo-PE), unsaturated phosphatidylethanolamine (DOPE), phosphatidic acid (PA), tetramyristoyl CL and dimyristoyl CL (BMP). The assay was performed by incubating purified PlaF in DDM micelles with the lipid substrates, followed by quantifying released FA by NEFA assay. The results are the mean ± SD of three independent experiments. C) Mechanism suggesting how PlaF-catalysed modifications of PLs may be linked to the pathways responsible for rapid response to environmental changes and *de novo* PL synthesis. Symbols of relevant lipids are shown in the upper right inset.

Cti and Cfa are peripheral cytoplasmic membrane proteins that catalyse the conversion of cis-unsaturated fatty acids (cis-UFA) to trans-UFA and cyclopropane FA in PLs, respectively. Such PLs with modified acyl chains change the membrane’s biophysical properties, which leads to the alteration of membrane protein function and finally affects cell physiology [84–86]. Another lipid homeostasis protein accumulated in Δ*plaF* is cytoplasmic membrane-bound PlsX, which initiates *de novo* PL biosynthesis by catalysing the first reaction in the synthesis of lysophosphatidic acid (LPA), a precursor of phosphatidic acid (PA), which is used for the synthesis of all PLs [87, 88]. Interestingly, no other proteins involved in the modification and turnover of PL or *de novo* PL, fatty acid and head group biosynthesis were significantly differentially abundant in Δ*plaF* (Table S9).

We suggest that PlaF might be responsible for the degradation of Cfa and Cti synthetized PLs after, a so far unknown, environmental signal diminishes. Here, we strengthen this hypothesis by demonstrating that purified PlaF shows *in vitro* hydrolytic activity towards PLs containing a cyclopropane-FA or unsaturated-FA (Fig. 5B). Furthermore, we show that purified PlaF catalyses the hydrolysis of PA (Fig. 5B). We analysed whether PlaF can hydrolyse cardiolipin (CL), which has biological functions in the modulation of bacterial membrane protein function and the assembly of membrane complexes through interaction with proteins, in *P. aeruginosa* and other bacteria [6, 8, 43, 89–91]. Purified PlaF released fatty acids from tetramyristoyl CL at a rate comparable to PL hydrolysis (Fig. 5B). Interestingly, the hydrolysis of dimyristoyl cardiolipin (BMP) by PlaF was eight times faster than that by tetramyristoyl CL. These results indicate that PlaF may be involved in remodelling the acyl chain composition of CL; however, it remains unknown whether PlaF-mediated cardiolipin degradation has biological relevance and if it is coupled to its biosynthesis because cardiolipin synthesis proteins were not identified by our proteomic analysis.

## Discussion

Genetic studies of bacteria with disrupted lipid biosynthesis pathways suggested phospholipids as critical determinants of bacterial envelope biogenesis [27, 92], which is not surprising considering the function of lipids for membrane protein stabilization, activation, complex assembly and cellular localization [93]. From recent integrative biology studies it is apparent that the bacterial cell cycle depends on the dynamic alteration of membrane structure achieved through biosynthesis, modification and turnover of proteins and lipids [94, 95]. Among these processes in bacteria, the degradation of membrane PLs is probably the least understood, with a major reason being the small number of known intracellular phospholipases [37, 38, 96]. We recently showed that the cytoplasmic membrane-bound phospholipase PlaF of *P. aeruginosa* modulates membrane PL composition, likely through the degradation of only a handful of PL species [38]. Here, we established the function of PlaF in biofilm biogenesis (Fig. 1) and swimming motility (Fig. 3B), strengthening its role in virulence that was previously determined in *Galleria mellonella* and bone marrow-derived macrophages [38]. We hypothesized that the PlaF-mediated alteration of membrane PL composition might indirectly cause adaptive changes to the biofilm-related proteome of *P. aeruginosa*.

### *Pathways and proteins affected in Δ*plaF

By focusing on identifying PlaF-regulated pathways, we performed quantitative proteomics of Δ*plaF* and WT biofilms. Strikingly, the results revealed the pleiotropic effect of PlaF in *P. aeruginosa* biofilm, as nearly ~8% of all *P. aeruginosa* PA01 proteins were differentially abundant (> 1.5-fold change) in Δ*plaF* under the chosen criteria. The importance of the regulatory function of PlaF is illustrated by the fact that the best-studied regulatory network of *P. aeruginosa*, the quorum sensing network, contain 627 genes or 11% (> 1.5-fold change determined by transcriptomics) of the *P. aeruginosa* genome [54]. However, we could not observe an obvious relationship between QS and PlaF networks, as none of the major QS regulators or regulated proteins was differentially abundant in Δ*plaF*, and most of the 22 overlapping Δ*plaF vs*. QS proteins/genes showed only a weak change (< 2-fold) in QS transcriptomic studies (Fig. S5). Proteomics results showed that the most strongly affected pathways in Δ*plaF* were, pyoverdine-mediated iron acquisition (Fig. 4), PprAB TCS-regulated biofilm formation and attachment (Fig. 2), and the proteolytic degradation system (Figs. 3C–3D, Scheme 1). The effect of PlaF on these pathways was verified by showing that Δ*plaF*, in comparison to WT, produces less pyoverdine (Fig. 4C), has lower intracellular iron concentrations (Fig. 4D), shows transcriptional upregulation of PprB-regulated genes (Fig. 2B), and exhibits less intracellular protease activity (Fig. 3E).

**Scheme 1:**
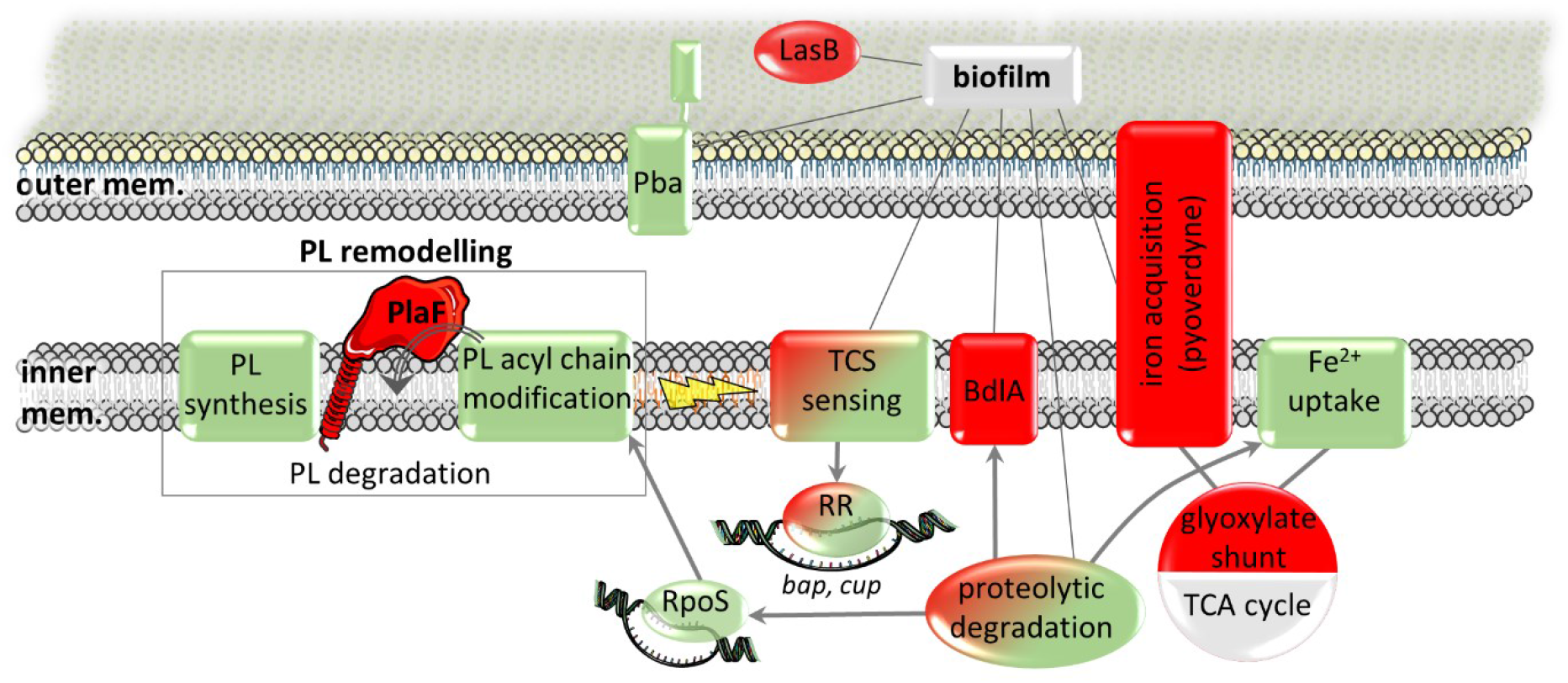
A model proposing the interplay between PlaF, PL remodelling and biofilm pathways. PlaF-catalysed PL degradation, Cti and Cfa catalysed modification of acyl chain of PLs and PlsX-pathway for PL synthesis jointly remodel (Fig. 5C) PL composition of inner membranes. Altered inner membrane PLs composition affects various biofilm-related proteins and pathways: signal sensing (TCS, BdlA) and transcriptional response (RR, RpoS), intracellular and extracellular (LasB) proteolytic degradation, peptidoglycan binding (Pba), iron acquisition (pyoverdine dependent and independent) and metabolism (GS and TCA). Grey arrows and lines show interaction demonstrated in this study and in the literature, respectively. Green and red colors indicate whether the system was more or less abundant (dual coloration indicates both tendencies) in the proteome of Δ*plaF*. Black lines indicate pathways and proteins with previously described link to biofilm.

### *How can PlaF simultaneously affect a large portion of the* P. aeruginosa *proteome?*

One of the possible answers to this question is the observation that PlaF affects several TCSs, thereby changing the transcriptional profile of *P. aeruginosa*. The relationship between PlaF and the regulation of gene expression is even more complex, keeping in mind that the alternative sigma factor RpoS and anti-anti-sigma factor HsbA were more abundant in Δ*plaF*. Bacteria widely use TCSs to modify gene expression in response to external cues [97, 98]. *P. aeruginosa* contains more than sixty TCSs regulating pathoadaptation in chronic and acute infections [57], and PlaF affected 14 TCSs that regulate the transcription of virulence genes involved in cell motility, adhesion, iron sensing and biofilm formation [57]. These TCS-regulated processes often require a defined membrane PL composition, as underlined by the observation that membrane PLs with polyunsaturated acyl chains reduce the flagellar adhesion of *E. coli* [99], PLs with saturated acyl chains promote the switch of *P. aeruginosa* from planktonic to the biofilm lifestyle [100], PLs with shorter acyl chains accumulate in older biofilms [22], and the synthesis of cyclopropanated PLs is coupled with the transition of *P. aeruginosa* in the stationary growth phase [24]. Adjustments of the membrane PL content during infection processes was observed not only in *P. aeruginosa* and evolutionary-related *E. coli* but also in *Salmonella* Typhimurium, *Staphylococcus aureus, Listeria monocytogenes* [100, 101] and *Mycobacterium tuberculosis* [25]; therefore, this adaptive mechanism is likely widespread among bacterial pathogens. It is worth emphasizing that little is known about the role of intracellular PLA-catalysed PL degradation in the modulation of membrane PL content and virulence adaptation in other bacteria, and further research is necessary to elucidate these aspects.

Future studies will reveal whether the environmental signals sensed by *P. aeruginosa* trigger the remodelling of membrane PLs by PlaF followed by transcriptional reprogramming of *P. aeruginosa* or whether PlaF-mediated PL remodelling is a consequence of altered gene transcription.

### How could PlaF affect TCS sensing and transduction?

TCS SKs are good targets for regulation through interaction with PLs, as these interactions provoke conformational changes in SKs or influence homodimerization of SKs that are associated with their ability to sense and transduce the signal [102, 103]. Despite comprehensive studies on TCSs based on their biological importance, little is known about the function and structure of transmembrane helical segments, and the functional role of PL-SK interactions is even less evident [102, 103]. To date, the only examples of TCS SKs regulated by PL are *Sinorhizobium meliloti* ExoS and *E. coli* DesK, whose activation/inactivation was shown to be regulated through direct interactions with PC or unsaturated PLs, respectively [10, 104]. Therefore, it is possible that the function of PlaF-affected TCS SKs may partially be modulated by the nonnative PL composition of Δ*plaF* membranes. Furthermore, some TCS that are differentially abundant in Δ*plaF* may be regulated through direct contact with the TM helix or soluble domain of PlaF, as a similar regulatory role of protein-protein interactions was shown for the WalI-WalK TCS [105] and PbgA-regulated PhoP-PhoQ TCS [106].

### PlaF-dependent TCS-mediated response in biofilm

The observation that PlaF affects TCSs which sense PL building blocks (phosphate by PhoR and fatty acids by TctD) provides a link between these TCS and the suggested function of PlaF for PL remodelling. This is in agreement with recent results demonstrating that *P. aeruginosa* adapts to phosphate limitation through membrane lipid remodelling [40]. Furthermore, the results demonstrating that six PlaF-dependent TCSs (RocA1, PhoR, FleR, PA4396, TctD, and PprA-PprB) regulate biofilm formation and adhesion [57, 61] are in agreement with the suggested role of PlaF in the indirect regulation of biofilm biogenesis through TCS-controlled gene transcription. Such a role for PlaF is strengthened by the finding that PlaF affects ten TCSs active in chronic infections tightly linked to a sessile lifestyle [107].

We observed biofilm-regulating PprAB TCS [50, 55, 108], which is highly abundant in Δ*plaF* biofilms (Fig. 2B). This TCS has been previously shown to activate the expression of fimbrial genes encoded in *tad, bap* and *cup* loci [55]. Here, we show by proteomics that biofilm-impaired Δ*plaF* overproduces cell surface appendages after 24 h, although they are important for the early stages of biofilm development [50]. Drastically weaker attachment of Δ*plaF* after 24 h was also observed by the CLSM analysis (Fig. 1B). These delayed PprAB responses in Δ*plaF* may explain why Δ*plaF* produces 3 times less biofilm than the WT after 24 h, while the levels of these two strains are balanced later in development (Fig. 1A). Such improper temporal coordination of the PprA-PprB response in Δ*plaF* may be a consequence of activity of altered PprA SK within the Δ*plaF* membrane with nonnative PL composition. Altogether, our data suggest an adaptive role of PlaF in biofilm formation, which agrees with previous observations that the biofilm adaptation of *P. aeruginosa* requires a fine-tuned membrane PL composition [22, 100].

It seems that the overproduction of SKs is a general response of Δ*plaF*, as six of eight differentially abundant SKs show higher abundance in Δ*plaF*. However, a higher abundance of phosphorylated RRs was not observed. After sensing the signal, SK phosphorylates RR, thereby changing its affinity to bind DNA, which finally results in altered gene transcription [109]. Furthermore, in addition to phosphorylation, SK bifunctional proteins can modulate gene expression by dephosphorylating RRs. This feedback mechanism involving modulation of the ability of SKs to regulate the concentration of phosphorylated RRs plays a critical role in the TCS-mediated response [109]. While most RRs are activated by phosphorylation [109], unphosphorylated CsgD of *Salmonella enterica* [110] and AgrR of *Cupriavidus metallidurans* are the active forms [111]. Despite a nearly 19-fold higher abundance of PprA in Δ*plaF*, we have not observed an increased abundance of phosphorylated PprB, yet transcription of *bap* and *cup* was upregulated, while that for *pqs* and *tad* from the *pprB* regulon was not upregulated. This suggests that phosphorylated and dephosphorylated forms of PprB may regulate the expression of a different set of genes. Although the regulatory mechanism of SKs by PlaF remains to be determined, it is tempting to speculate that PlaF-mediated PL remodelling or direct PlaF-SK interactions modulate phosphatase or kinase activity of PprA and other PlaF-dependent SKs, thereby activating the regulator to a different extent, which is important for coregulated expression of functionally related genes. Such a role of PlaF would explain the inability of Δ*plaF* to coordinate complex processes such as biofilm formation and infection.

One of the feedback mechanisms of TCS is an alteration of the amount of RR by proteolytic degradation. Despite proteomics results revealing a differential abundance of several known and putative proteases (Fig. 3A) and semitryptic peptides (Table S7) in Δ*plaF*, we did not observe proteolytic degradation of PlaF-dependent TCS proteins which might be further degraded to amino acids [112]. Nevertheless, we identified semitryptic peptides generated by proteolysis of several other TCS proteins (CbrA, CbrB, PhoP, PA4380, and PA1397), thereby confirming the turnover of TCS proteins in Δ*plaF*. Here identified, significantly different levels of abundance of semitryptic peptides of 45 proteins in Δ*plaF* (Table S7) confirmed the indirect role of PlaF on the protein turnover system, which might be one of the mechanisms by which PlaF affects biofilm biogenesis, which relies on proteolytic degradation [113, 114]. For example, attenuated virulence and reduced biofilm formation in Δ*plaF* might partly be due to the lower abundance of a well-known virulence and biofilm factor, an extracellular protease LasB, in this strain [115]. Significantly different levels of abundance of semitryptic peptides of sigma factor RpoS, which regulates the transcription of biofilm genes [116], and sensor protein BdlA, which modulates the activity of enzymes regulating c-di-GMP signalling, in Δ*plaF* further strengthen the indirect regulation of biofilms through modulation of gene transcription by PlaF-dependent protein degradation.

### Relationship between PlaF, iron acquisition and metabolism

The interdependence of iron acquisition, biofilm formation and virulence is well documented in *P. aeruginosa* [56, 75, 117] and other major pathogens [25, 118, 119]. However, understanding the relationship between iron acquisition and membrane PL composition in bacteria is limited. To our knowledge, only one recently published report shows downregulation of the iron uptake pathway in an *S. meliloti* mutant deficient in the synthesis of phosphatidylethanolamine [104]. Our proteomics results revealed downregulated pyoverdine-mediated iron uptake in Δ*plaF* biofilms, while pathways that rely on low-affinity iron-chelating siderophores, pyochelin and citrate, and siderophore-independent phenazine-assisted Fe^2+^ uptake were upregulated [69, 79, 80]. This is in agreement with the lower concentrations of pyoverdine in the supernatant and iron inside the cells of Δ*plaF* (Figs. 4C and 4D). Synthesis of pyoverdine is activated under iron limitation by the ferric uptake regulator (Fur) [120, 121], which was not detected in the proteomic study of Δ*plaF*, although it is likely abundant at low concentrations because the products of Fur-regulated genes were detected. A low Fur concentration may explain why the decrease in intracellular iron concentrations in Δ*plaF* did not induce the expression of pyoverdine synthesis enzymes.

We observed that pyoverdine limitation in Δ*plaF* results in the upregulation of pyochelin synthesis enzymes PchA and PchC but not the transport of pyochelin across the inner membrane by FptX and outer membrane by receptor FptA, as the abundance of both transporters was not affected in Δ*plaF*. In addition to pyoverdine- and pyochelin-mediated Fe^3+^ uptake, *P. aeruginosa* has developed the ability to acquire Fe^2+^ *via* the TonB-independent citrate and Feo systems, whose inner membrane transporters, MgtA [69] and FeoB [122], were more abundant in Δ*plaF*, respectively. This, together with a higher abundance of PhnA, involved in the synthesis of the redox-active molecule pyocyanin, which promotes Fe^2+^ acquisition, suggests the predominant acquisition of Fe^2+^ in Δ*plaF*. These data are consistent with the increased importance of citrate-mediated iron acquisition in the biofilm of the pyoverdine synthesis mutant (Δ*pvdA*) [123] and upregulation of the Feo iron acquisition system under low-iron conditions of a static biofilm [124, 125]. Our results show an important virulence function of PlaF in modulating iron acquisition under static biofilm conditions; however, it remains to be shown whether PlaF exerts a similar function during chronic infections strongly dependent on iron acquisition and biofilm formation [124]. The link between iron acquisition and PlaF goes beyond protein abundance, as the activities of other inner membrane-associated iron acquisition proteins, e.g., pyoverdine synthesis monooxygenase PvdA [126] or transporter FeoB, may be modulated by the nonnative PL composition of Δ*plaF* or phosphorylation, as was shown for MgtA, whose activity is regulated by cardiolipin and phosphorylation [6].

Iron is not indispensable only for biofilms but also for growth and primary metabolism as a cofactor of many metabolically important enzymes, e.g., enzymes from the electron-transport chain as well as tricarboxylic acid (TCA) cycle [81]; therefore, bacteria accumulates iron in the lag phase [82]. This agrees with our observation that planktonic Δ*plaF* culture has a prolonged lag phase (Fig. 4E) and decreased ability to take up iron (Fig. 4C). We observed that PlaF affects primary metabolism through its influence on the partitioning carbon flux between the TCA cycle and GS. This is achieved by the differential abundance of phosphorylated isocitrate dehydrogenase (TCA pathway) and isocitrate lyase (GS pathway) in Δ*plaF*, which leads to the downregulation of GS. The effect of PlaF on GS may be a further mechanism by which PlaF affects iron homeostasis, as it was previously shown that *P. aeruginosa* cells lacking a GS pathway have dysregulated iron homeostasis [81, 127]. Positioning at the intersection of energy and iron homeostasis [81], GS is a pathway required for *in vivo* infections by *P. aeruginosa* [128], *M. tuberculosis* [129] and even fungal pathogens [130, 131].

A prolonged lag phase, an altered GS pathway, RpoS and TCS-mediated signalling, and membrane PL composition are typical stress responses of bacteria [27, 116, 132–134]. Deletion of *plaF* gene in *P. aeruginosa* has unforeseen consequences for all these processes which indicates that PlaF might coordinate the adaptive stress response by a novel mechanism involving modulation of the cytoplasmic membrane PL composition or interactions with other proteins.

### Relationship between PlaF and phospholipid homeostasis

Increasing evidence of protein-lipid interactions [6, 7, 13, 93] and genetic studies [27, 92, 104] have suggested that bacteria use distinct PL molecules to modulate specific cellular pathways. This concept raises the question of how bacteria specifically remove PL signalling molecules to terminate or modulate regulated pathways. Simple dilution of PL modulators during cell division seems inappropriate, as it is slow or even impossible when growth is inhibited. Another possibility is a system for rapid responses to environmental changes, which relies on altering the PL composition under stress conditions [83, 135]. This system involves modifications of the PL structure by Cti-catalysed *cis-trans* isomerization of unsaturated PL acyl chains and Cfa-catalysed conversion of the PL double bond to a cyclopropyl group [83, 136]. Cti and Cfa belong to this rapid response system and have a particularly important function under stress conditions [83, 135].

We observed that the abundance of Cfa and Cti increased in Δ*plaF* (Fig. 5A); however, Cfa-produced cyclopropanated PLs did not accumulate in Δ*plaF* [38]. As Cti and Cfa cannot catalyse reversible reactions, another mechanism for converting the cyclopropyl group and trans-double bond to the cis-configuration of the acyl chain in PLs is unknown [83]; however, it is likely that *P. aeruginosa* degrades these PLs after the stimulus is removed. Our biochemical results show that PlaF can hydrolyse cycloproanated and unsaturated PLs (Fig. 5B); therefore, we suggest that the PL degrading function of PlaF and PL acyl chain modifying functions of Cti and Cfa are responsible for maintaining the acyl chain composition of membrane PLs. This is in agreement with a model by Witholt and Pedrota (1999), who proposed that Cti-catalysed isomerization requires the previous release of fatty acids from PLs by phospholipase A, as Cti does not show activity on pure PLs but shows activity on PLs isolated from bacterial cells [137].

Although it needs to be experimentally determined if PlaF competes with Cfa and Cti for the same substrate *in vivo*, their functional relationship in the cell is underlined by several biochemical and physiological properties that they share: a) They are colocalized at the cytoplasmic membrane where they catalyse the conversion of PLs [38, 83, 136]; b) PlaF and Cfa have been shown to be regulated through dimerization; c) FA products have been shown to regulate PlaF activity and dimeric state [38], while trans-configurations in acyl chains of PLs seem to regulate the membrane insertion of Cti, thereby inhibiting isomerization reaction [85]; d) PlaF and Cti are constitutively expressed at low levels, while the expression of Cfa begins when the cells enter stationary phase; e) PlaF [38] and *E. coli* Cfa [138] show a preference for the *sn*-1 position of PLs (regiospecificity of Cti is unknown); and f) RpoS and Fe^3+^, both linked to PlaF, respectively activate the transcription of Cfa and the enzymatic activity of Cti [136, 139].

Our lipidomic study [38] showed that the absence of PlaF in *P. aeruginosa* resulted in the accumulation of some PL species, while several other PL species were depleted, thus indicating altered PL biosynthesis. Indeed, proteomics revealed a higher abundance of PlsX, a key enzyme of PL biosynthesis, which catalyses the synthesis of LPA; LPA is subsequently converted to PA, which serves as the substrate for the synthesis of all PLs [87]. The membrane insertion, and thus access to the substrate, of the peripheral membrane protein PlsX is regulated through membrane fluidity [83, 140]; therefore, it can be affected by the PlaF-Cti-Cfa system. Furthermore, as PlaF shows hydrolytic activity with PA (Fig. 5B), it can interfere with the PlsX pathway by regulating product concentrations.

Based on lipidomics [38] and proteomics results, we suggest a model (Fig. 5C) in which deletion of the *plaF* gene in *P. aeruginosa* leads to higher amounts of Cfa and Cti and subsequent enrichment of PLs containing trans-UFA and cyclopropane-FA. Such a nonnative membrane with affected fluidity triggers the binding of PlsX to the membrane and upregulation of PlsX production, which in turn enhances *de novo* PL synthesis to maintain the balance of unmodified and modified PL, which is essential for membrane homeostasis.

## Methods

### Liquid chromatography coupled tandem mass spectrometric-based proteomic analysis

Five cultures of *P. aeruginosa* PAO1 (wild-type, WT) and Δ*plaF* (mutant) inoculated from a single colony grown on an LB agar plate were cultivated overnight at 37°C in LB medium [141] under aeration. These cultures were diluted with LB medium to an optical density (OD_580nm_) of 0.1, and 100 μl of each culture was transferred to 16 wells of a 96-well microtiter plate (MTP). Biofilm was grown for 24 h as described previously [38], and the samples for mass spectrometric (MS) analysis were prepared by harvesting the cells (21,000 *g*, 4 °C, 30 min) from 16 wells for each of 10 cultures (5 biological replicates of *P. aeruginosa* WT and Δ*plaF)*, followed by the removal of supernatant and immediate suspension in lysis buffer (0.03 M Tris-HCl, 2 M thiourea, 7 M urea, pH 8.5) to prevent proteolysis.

Protein lysates were further prepared for MS analysis [142]. Briefly, proteins were shortly separated in an acrylamide gel, stained with Coomassie brilliant blue, reduced with dithiothreitol, alkylated with iodoacetamide and digested with trypsin overnight. The resulting peptides were extracted from the gel, dried in a vacuum concentrator, and dissolved in 0.1 % trifluoroacetic acid (TFA). About 500 ng of peptides were separated using an UltiMate 3000 RSLCnano chromatography system (Thermo Fisher Scientific). First, peptides were concentrated for 10 minutes at a flow rate of 6 μl/min with 0.1 % (v/v) TFA as mobile phase on a trap column (Acclaim PepMap100, 2 cm length, 3 μm C18 particle size, 100 Å pore size, 75 μm inner diameter, Thermo Fisher Scientific) and finally separated at 60 °C using a 2 h gradient from 4 to 40 % (v/v) solvent B (solvent A: 0.1% (v/v) formic acid in the water, solvent B: 0.1 % (v/v) formic acid, 84 % (v/v) acetonitrile in water) at a flow rate of 300 nl/min on an analytical column (Acclaim PepMapRSLC, 25 cm length, 2 μm C18 particle size, 100 Å pore size, 75 μm inner diameter, Thermo Fisher Scientific). Separated peptides were directly injected in an online coupled QExactive plus mass spectrometer (Thermo Fisher Scientific) using an electrospray ionization nano source and distally coated SilicaTip emitters (New Objective, Woburn, MA, USA) at a spray voltage of 1.4 kV. The mass spectrometer was operated in data-dependent positive mode using a capillary temperature of 250 °C.

Firstly, full scans were recorded in the orbitrap in profile mode at a resolution of 70,000 over a scan range from 350 to 2000 m/z (maximal ion time 80 ms, automatic gain control 3,000,000). Secondly, up to ten two and threefold charged precursor ions were sequentially selected within a 2 m/z isolation window by the quadrupole, fragmented by higher-energy collisional dissociation and fragments analyzed in the orbitrap (resolution 17,500, 60 ms maximal ion time, automatic gain control 100,000, available scan range 200 to 2000 m/z). Fragment spectra were recorded in centroid mode, and already fragmented precursors were excluded from analysis for the next 100 s.

After data acquisition, spectra were subjected to peptide and protein identification using the MS Amanda search engine triggered by proteome discover (version 1.4., Thermo Scientific). Here, 5582 *P. aeruginosa* PAO1 protein sequences obtained from Pseudomonas Genome Database (PGD, http://www.pseudomonas.com) [143] were *in silico* cleaved by trypsin cleavage specificity (up to two missed cleavage sites), methionine oxidation, deamidation of asparagine and glutamine and N-terminal acetylation as variable and carbamidomethylation as fixed modification considered. Mass tolerances of 5 ppm were used for precursor ions and 0.02 Da for fragment ions. The percolator node was used for identification validation (false discovery rate on peptide and protein level 1%); only proteins were considered to be identified, showing at least two different peptides. The MS proteomics data have been deposited to the ProteomeXchange Consortium via the PRIDE [144] partner repository with the dataset identifier PXD037737.

Subsequently, a quantitative analysis based on precursor ion intensities was carried out by the Progenesis IQ for proteomics software (version 2.0.5387, Nonlinear Dynamics) and linked to the previously identified peptides and proteins. Sum intensities of non-ambiguous features belonging to certain protein groups were used for further calculations. Here, log-transformed data were used for calculating Student’s t-tests for proteins with valid values in all samples. As we focused this analysis instead on a global view of PlaF-related changes instead of concentrating on single proteins, we did not correct for multiple testing but considered only proteins as differentially abundant, showing a *p*-value ≤ 0.05 and a fold change Δ*plaF vs*. WT ≥ 1.5.

Furthermore, we mined the MS data for phosphorylated peptides. [142] Here, we used MaxQuant (version 1.6.1.0, MPI for Biochemistry, Planegg, Germany) for phosphorylated peptide identification and quantification with standard parameters if otherwise stated. PGD [143] entries (see above) were used for the search, carbamidomethylation configured as fixed and methionine oxidation, N-terminal acetylation and phosphorylation at serine, threonine and tyrosine as variable modifications. The ‘match between runs’ function was enabled to facilitate a transfer of identification information between the signals of the different runs. Peptides and proteins were identified at a false discovery rate of 1%, and *p*-values calculated with Student’s *t*-tests on log-transformed intensities of phosphorylated peptides. Furthermore, a search with semitryptic cleavage specificity was carried out using the same settings, except phosphorylation as variable modification.

### Functional assignment

*P. aeruginosa* proteins involved in iron homeostasis (*Pa*FeHo) and PL homeostasis (*Pa*PLHo) were obtained by a comprehensive literature search. In addition, a diamond BLASTp tool [145] from the PGD [143] was used to identify *P. aeruginosa* homologs of PL homeostasis enzymes from *E. coli*. An assignment of biological processes to *P. aeruginosa* proteins was performed with COG [46].

### Quantitative PCR

RNA was isolated with the NucleoSpin^®^ RNA preparation kit (Macherey–Nagel, Düren, Germany), and genomic DNA was quantitatively removed from RNA samples using RNase-Free DNase kit (Qiagen, Hilden, Germany) and Ambion^™^ DNA-free^™^ DNase kit (Thermo Scientific^™^, Darmstadt, Germany) according to the manufacturer’s recommendations. One μg of RNA was transcribed into cDNA using the Maxima First Strand cDNA Synthesis Kit (Thermo Scientific^™^, Darmstadt, Germany). For the qPCR, 50 ng of cDNA was mixed with SYBR Green/ROX qPCR Master mix (Thermo Scientific^™^, Darmstadt, Germany) to a total volume of 20 μl.

### Pyoverdine quantification

For the spectrophotometric quantification of pyoverdine at 400 nm wavelength, cell-free supernatant was obtained by filtration of the *P. aeruginosa* cultures, grown under biofilm conditions (see above) for 24 h, through a filter with 0.2 μm pore size. The extinction coefficient of pyoverdine of 19,000 M^-1^ cm^-1^ was used for calculations [87].

### Iron quantification

Quantification of iron in *P. aeruginosa* WT and Δ*plaF* cultures was done by a modification of the colorimetric method of Tamarit *et al*. (2006). [146] Cells of 24 h biofilm cultures were grown and harvested as described for proteomic analysis (see above) and washed with 1 ml water (Milli-Q). Cells were suspended in 400 μl HNO_3_ (3 % v/v) and incubated overnight at 98 °C. Insoluble residues were removed by centrifugation (10 min, 15,000 *g*), and 300 μl of supernatant was used for further analysis. A standard curve with FeSO_4_ (500 μM – 7.8 μM) was used for quantification.

### PlaF activity assay

PlaF was purified in the presence of n-dodecyl β-D-maltoside (DDM) as described previously [38]. Phospholipids and phosphatidic acid purchased from Avanti Polar Lipids (Alabaster, USA) were prepared for enzyme activity assays (25 μl enzyme + 25 μl substrate) as described previously [147]. The amount of released fatty acids by PlaF was determined using the NEFA-HR(2) kit (Wako Chemicals, Richmond, USA) [38].

### Protease assay

Protease activity was measured according to the manufacturer instructions of the Pierce Fluorescent Protease Assay Kit from Thermo Fisher Scientific. Briefly, *P. aeruginosa* WT and Δ*plaF* were cultivated under biofilm conditions described for proteomic analysis. Cells were harvested from 8 wells for each of 10 cultures (five biological replicates of *P. aeruginosa* WT and Δ*plaF)* followed by removal of supernatant and suspension in cold 500 μl TBS buffer (0.025 M Tris, 0.15 M NaCl, pH 7.2). Afterward, cells were sonicated three times for 20 seconds on ice. 100 μl of cell lysates were mixed with 100 μl of FTC-casein (10 μg/ml dissolved in TBS buffer). Fluorescence measurement was carried out immediately for 60 minutes at room temperature with standard fluorescein excitation and emission wavelengths of 485 nm and 538 nm, respectively. Trypsin served as standard and was used in a serial dilution (0.5 μg/ml - 0.0078 μg/ml).

### Fluorescence imaging of biofilm in flow chambers

*P. aeruginosa* WT and Δ*plaF* biofilms were grown on a microscope cover glass (24 mm x 50 mm, thickness 0.17 mm, Carl Roth GmbH & Co. KG, Karlsruhe, Germany), which was fixed with PRESIDENT The Original light body silicon (Coltène/Whaledent AG, Altstätten, Switzerland) on the upper side of the three-channel flow chambers [148]. The flow chambers and tubes (standard tubing, ID 0.8 mm, 1/16” and Tygon Standard R-3607, ID 1.02 mm; Cole-Parmer GmbH, Wertheim, Germany) were sterilized by flushing with sterile chlorine dioxide spray (Crystel TITANIUM, Tristel Solutions Ltd., Snailwell, Cambridgeshire, United Kingdom). Afterwards, the flow chambers were filled with 1 % (v/v) sodium hypochlorite, and the tubes were autoclaved. All biofilm experiments were performed at 37 °C with a ten-fold diluted LB medium. Before inoculation, the flow chamber was flushed with a 1:10 diluted LB medium for 30 minutes with a flow rate of 100 μl/min using the IPC12 High Precision Multichannel Dispenser (Cole-Parmer GmbH, Wertheim, Germany). For inoculation, an overnight culture of *P. aeruginosa* PAO1 or Δ*plaF* was adjusted to an OD_580nm_ of 0.5 in 1:10 diluted LB medium. The diluted culture (300 μl) was inoculated in each channel. After the interruption of medium supply for 1 h, the flow (50 μl/min) was resumed, and the biofilm structure was analyzed after 24, 72, and 144 h grown at 37 °C. For visualization, the cells were stained with propidium iodide and SYTO 9 dyes using the LIVE/DEAD^™^ BacLight^™^ Bacterial Viability Kit (Thermo Fisher Scientific). Imaging of biofilm was performed using the confocal laser scanning microscope (CLSM) Axio Observer.Z1/7 LSM 800 with airyscan (Carl Zeiss Microscopy GmbH, Germany) with the objective C-Apochromat 63x/1.20W Korr UV VisIR. The microscope settings were as described before [37]. The analysis of the CLSM images and three-dimensional reconstructions were done with the ZEN software (version 2.3, Carl Zeiss Microscopy GmbH, Germany). Quantification of mean thickness and height of biofilm was performed using the BiofilmQ software [149]. Experiments were repeated two times, each with one biological replicate that was analyzed at three different points by imaging a section of 100 x 100 μm.

## Acknowledgments

This study was funded by the Deutsche Forschungsgemeinschaft (DFG, German Research Foundation), project CRC 1208 (number 267205415) to FK and KEJ (subproject A02) and KS, GP and DWL (subproject Z01). We thank C. Strunk and R. Molitor (HHU Düsseldorf) for their help with phospholipase activity assay and swimming motility assay, respectively.

## Competing interests

The authors declare no competing interests.

## Supplementary material

**Figure S1:**
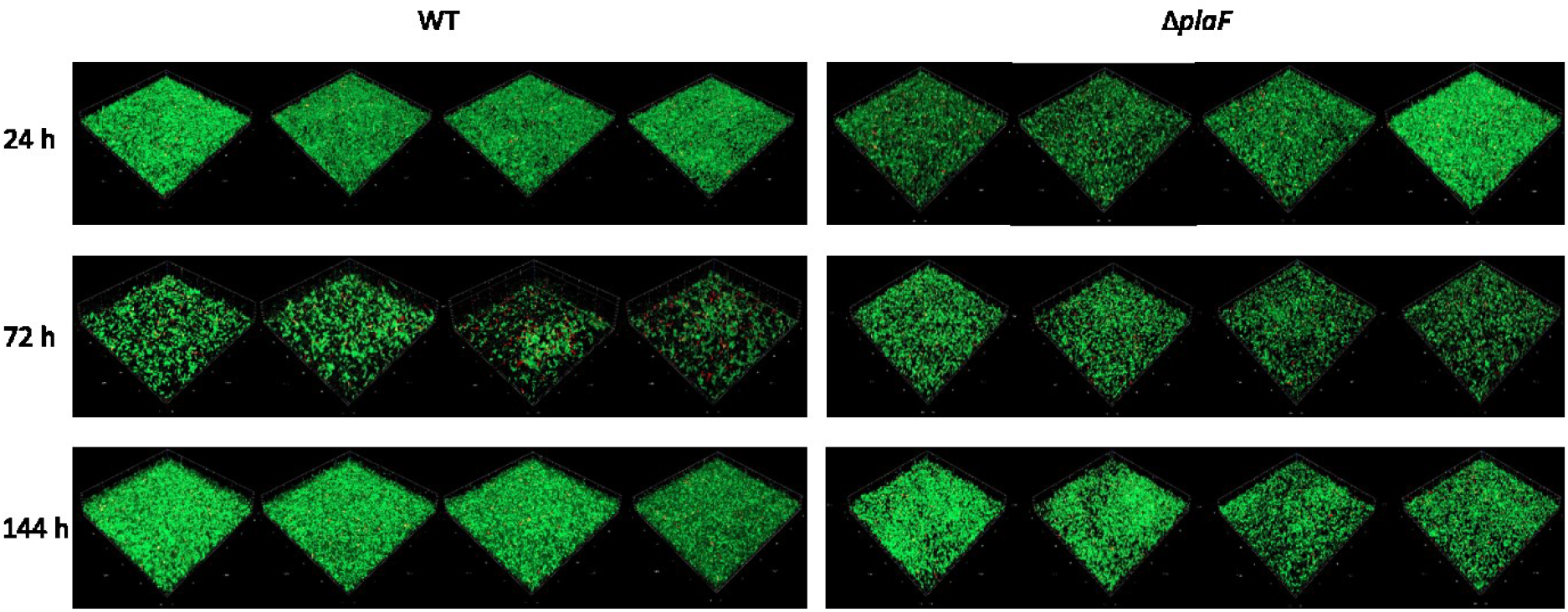
Biofilm architecture of Δ*plaF* and WT analysed by CLSM after 24, 72, and 144 h of growth at 37 °C in a flow cell with a continuous supply of LB medium. Images of two biological replicates obtained from two independent experiments.

**Figure S2:**
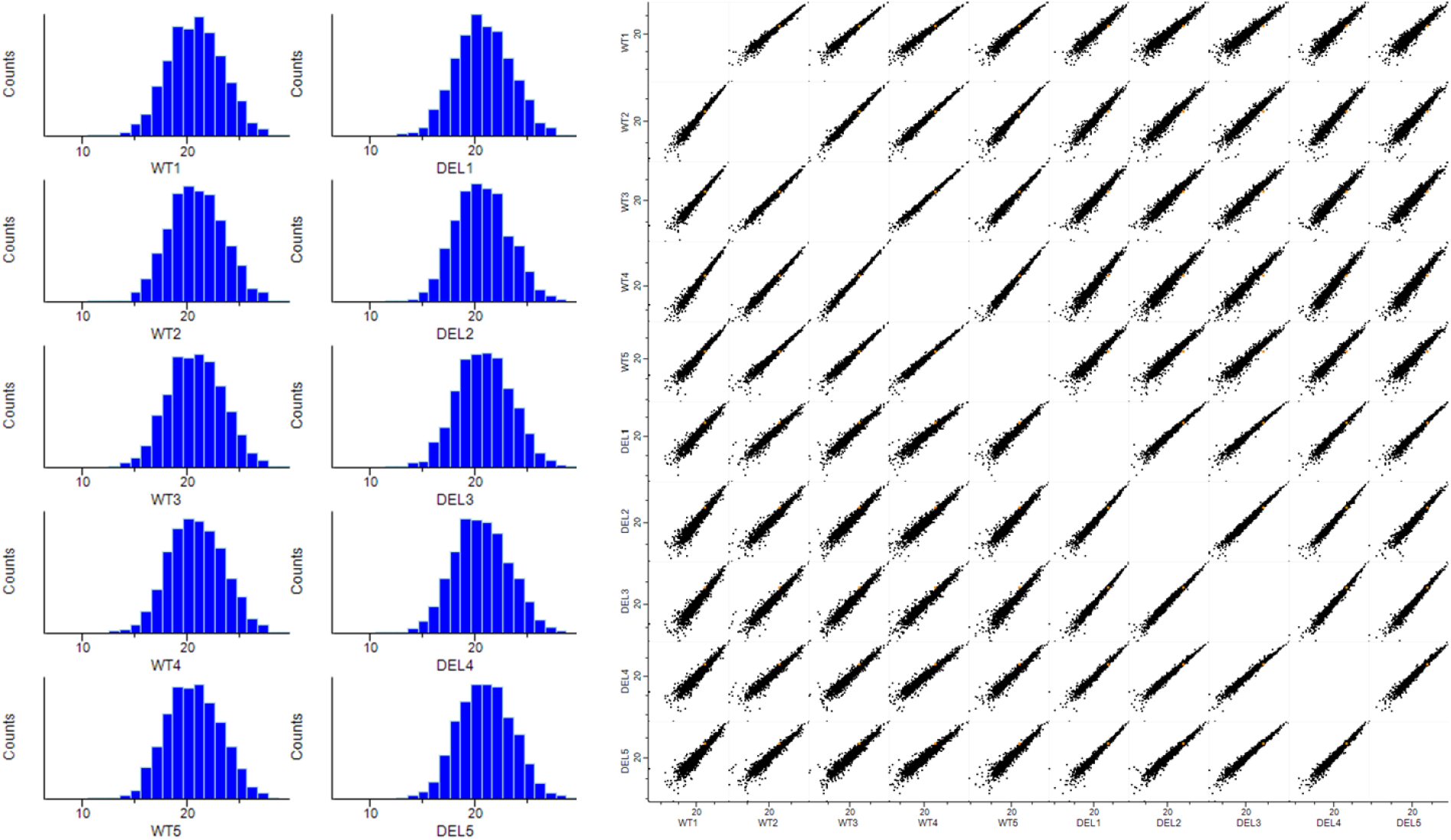
MS data quality. Quality control of quantitative MS data reveals comparable normal distributions, such as distributions of quantitative values. There is a correlation between normalized intensity values in scatter plots within the respective sample groups.

**Figure S3:**
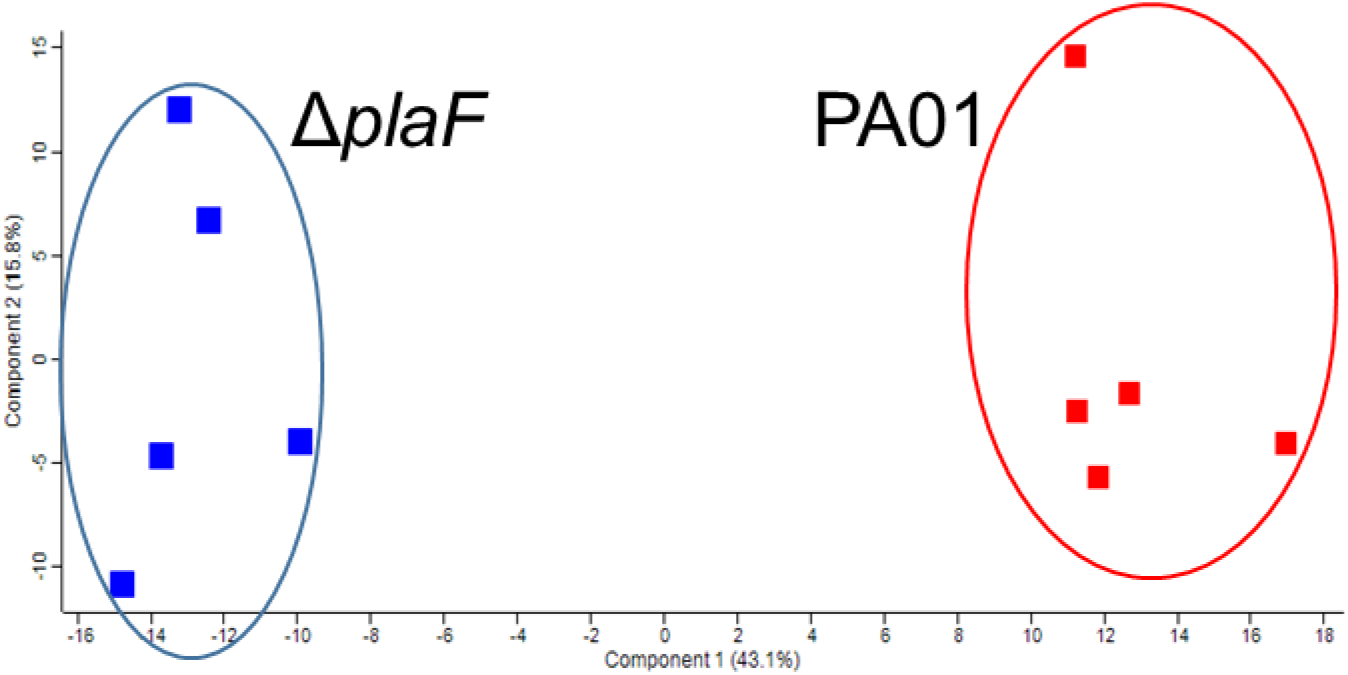
Proteome-wide comparison. Principal component analysis of normalized intensity values separated both sample groups fairly well based on component 1, which explained 43% of the observed variance.

**Figure S4:**
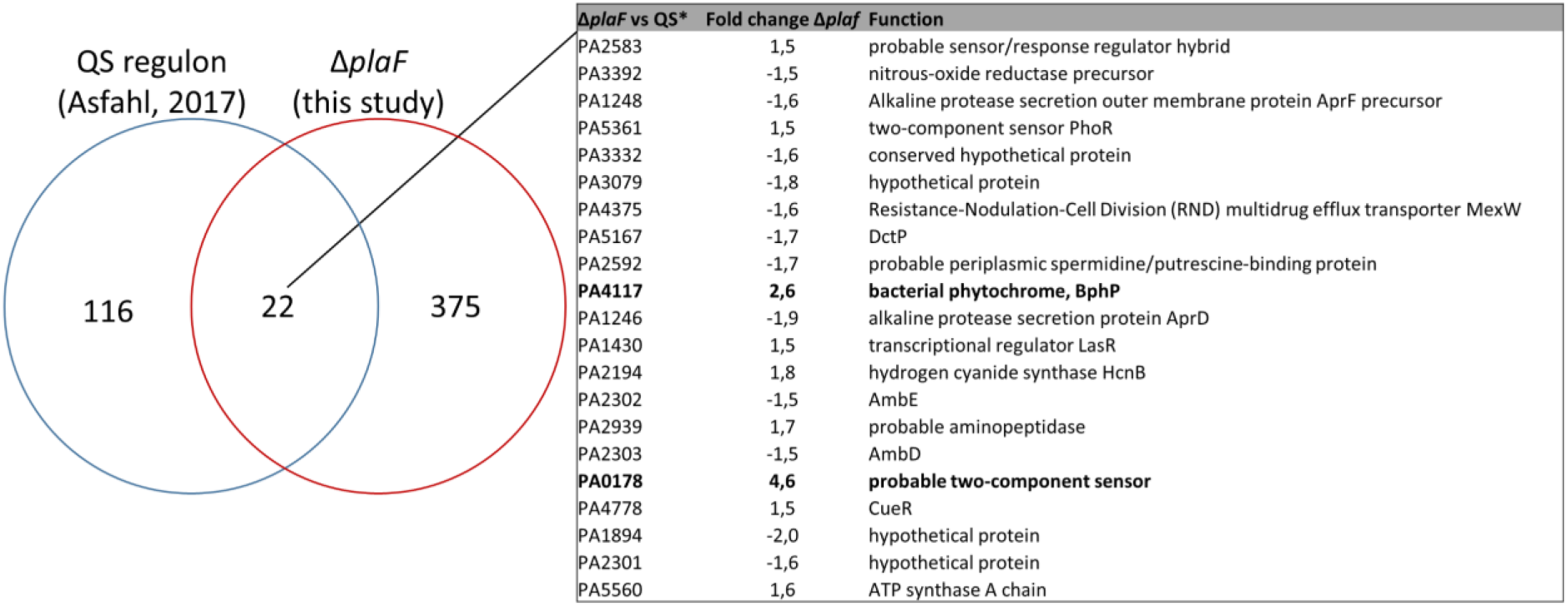
Proteins identified in Δ*plaF* (this study) overlapping with the QS regulon (*) represented by 139 gene products identified as differentially transcribed among the WT and isogenic Δ*lasR*Δ*rhlR* mutant [54]. Proteins less and more abundant in Δ*plaF* are indicated by negative and positive values, respectively. Locus tags and functions are from the PGD [143].

**Figure S5:**
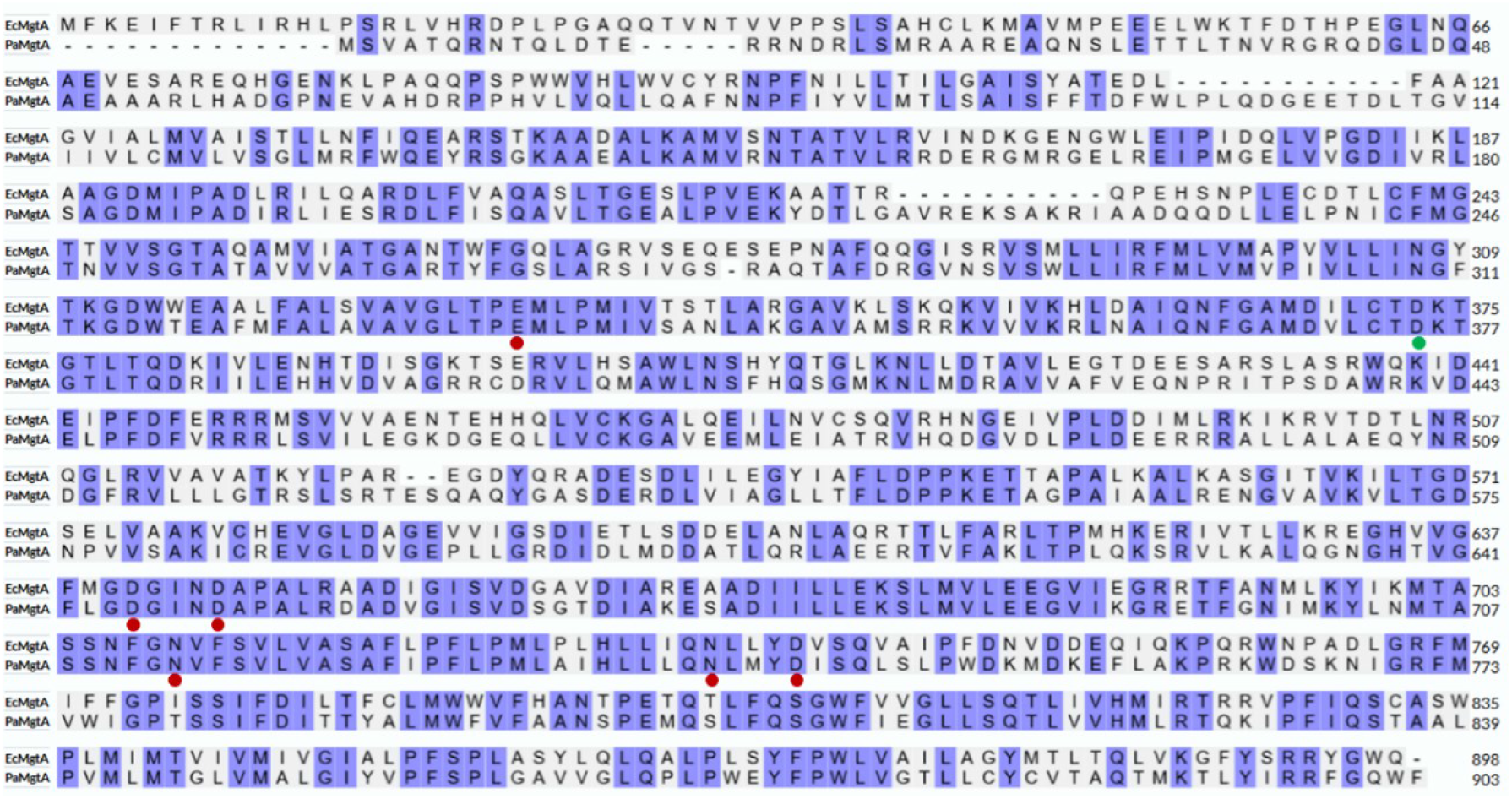
Sequence alignment of *P. aeruginosa* and *E. coli* MgtA proteins. Identical residues are highlighted in purple, and Mg^2+^ binding residues and phosphorylated D373 of *E. coli* MgtA are indicated by red and green dots, respectively.

**Figure S6:**
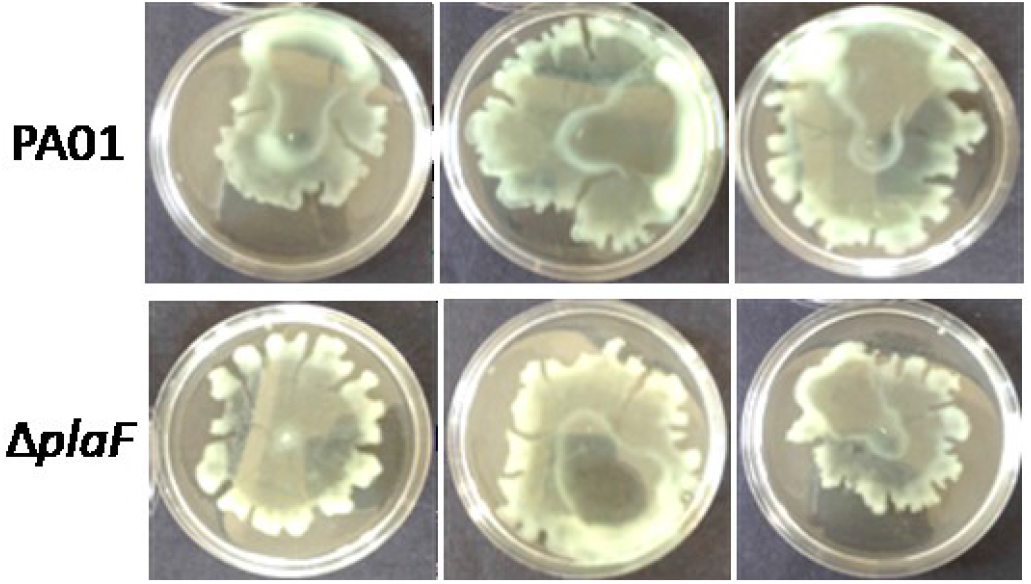
Twitching (A) and swarming (B) of Δ*plaF* were comparable to those of the WT after 24 h of incubation at 37 °C. Here, 3 representative replicates of 9 biological replicates are shown.

**Figure S7:**
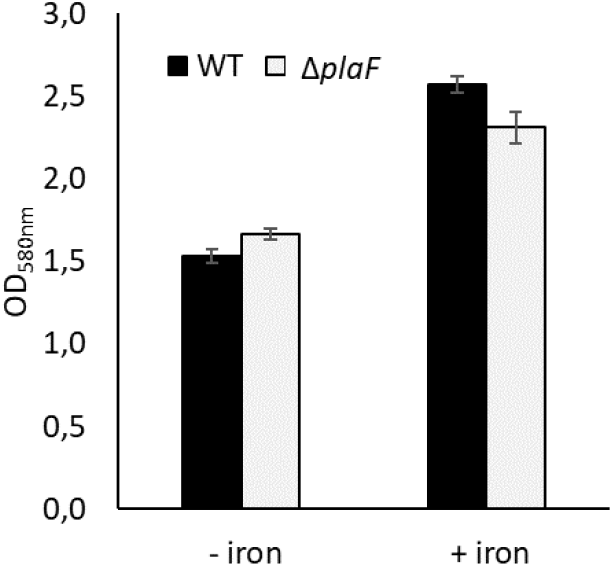
Growth of *P. aeruginosa* PA01 and Δ*plaF* in Erlenmayer flasks (aeration) for 24 h at 37 °C in M9 medium depleted in iron (-iron) or containing 100 μM FeSO_4_ (+ iron).

**Figure S8:**
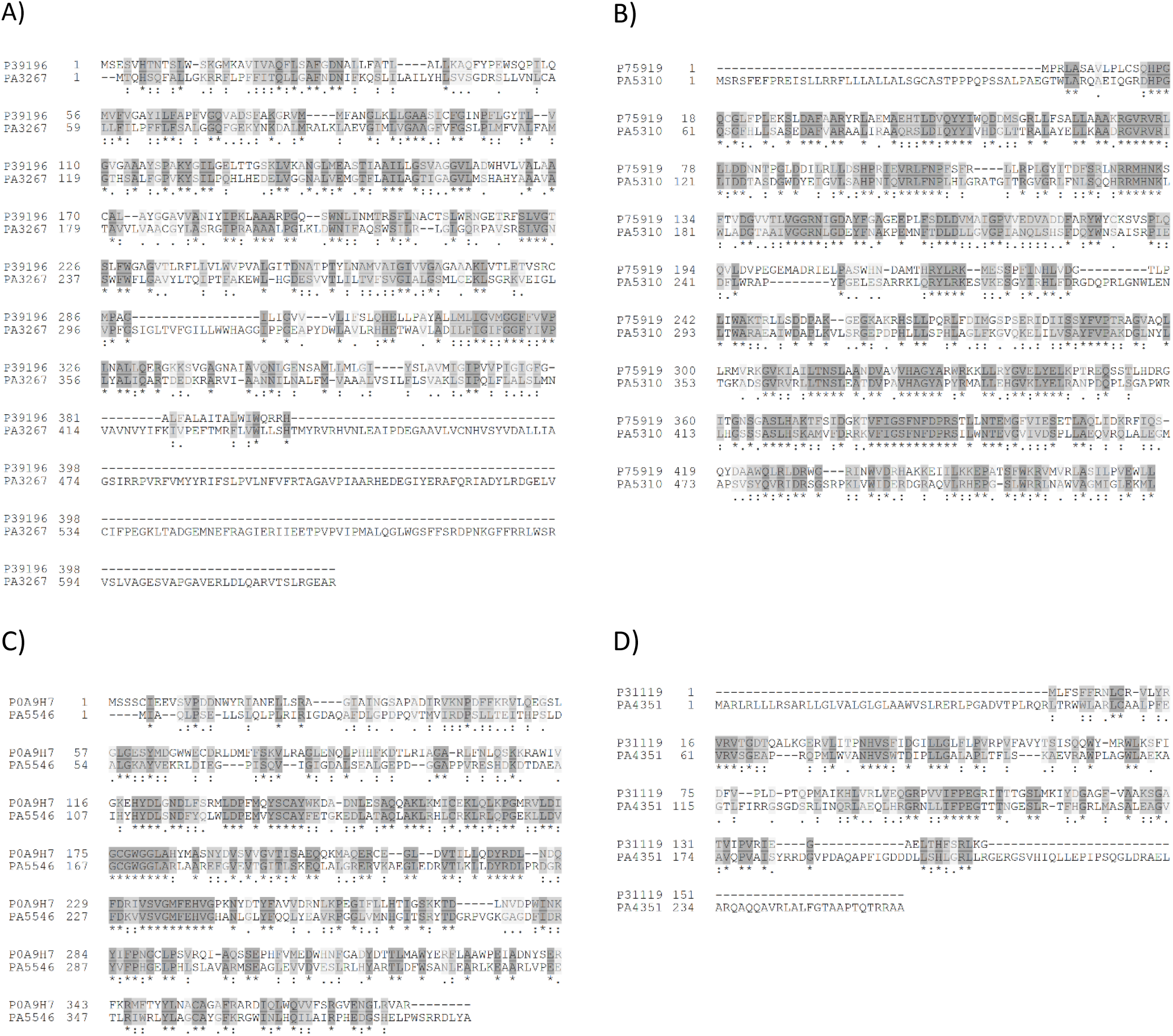
Identification of PL homeostasis proteins in *P. aeruginosa* by sequence alignment. The sequences of *E. coli* proteins were used for a BLAST search of the PGD [143], and the closest identified *P. aeruginosa* PA01 homologues were aligned with the respective *E. coli* proteins. A) *E. coli* LplT is 30.1% identical to *P. aeruginosa* PA3267. B) *E. coli* ClsC (P75919) is 38.4% identical to *P. aeruginosa* PA5310. C) *E. coli* Cfa (P0A9H7) is 41% identical to *P. aeruginosa* PA5546. D) The acyltransferase domain (residues 1-150) of *E. coli* Aas (P31119) is 31.4% identical to *P. aeruginosa* PA4351.

### Tables in supporting material are available upon request

**Table S1:** Proteins quantified in WT and Δ*plaF* by electrospray-ionization mass spectrometry.

**Table S2:** Proteins differentially abundant in WT and Δ*plaF*.

**Table S3:** Enrichment analysis of protein cellular localization in WT and Δ*plaF*.

**Table S4:** The functional classification of proteins differentially abundant in Δ*plaF*.

**Table S5:** Shared proteins identified in proteomics studies of Δ*plaF* (this study), attachment to glass and plastic surfaces [48], and biofilm formation [47].

**Table S6:** Phosphorylated proteins in WT and Δ*plaF* identified by MS analysis.

**Table S7:** Semitryptic peptides identified in Δ*plaF* and WT by proteomics.

**Table S8:** *P. aeruginosa* iron-homeostasis (*Pa*FeHo) network derived from the literature.

**Table S9:** Proteomics analysis of *P. aeruginosa* PA01 proteins involved in PL homeostasis.

## Notes

### Competing Interest Statement

The authors have declared no competing interest.

